# 3D single-molecule super-resolution imaging of microfabricated multiscale fractal substrates for self-referenced cell imaging

**DOI:** 10.1101/2023.11.07.566090

**Authors:** Clément Cabriel, R. Margoth Córdova-Castro, Erwin Berenschot, Amanda Dávila-Lezama, Kirsten Pondman, Séverine Le Gac, Niels Tas, Arturo Susarrey-Arce, Ignacio Izeddin

## Abstract

Microstructures arrayed over a substrate have shown increasing interest due to their ability to provide advanced 3D cellular models, which open new possibilities for cell culture, proliferation, and differentiation. Still, the mechanisms by which physical cues impact the cell phenotype are not fully understood, hence the necessity to interrogate cell behavior at the highest resolution. However, cell 3D high-resolution optical imaging on such microstructured substrates remains challenging due to their complexity, as well as axial calibration issues. In this work, we address this issue by leveraging the self-referenced characteristics of fractal-like structures, which simultaneously modulate cell growth and serve as axial calibration tools. To this end, we use multiscale 3D SiO_2_ substrates consisting of spatially arrayed octahedral features of a few micrometers to hundreds of nanometers. Through optimizations of both the structures and optical imaging conditions, we demonstrate the potential of these 3D multiscale structures as calibration tools for 3D super-resolution microscopy. We use their intrinsic multiscale and self-referenced nature to simultaneously perform lateral and axial calibrations in 3D single-molecule localization microscopy (SMLM) and assess imaging resolutions. We then utilize these substrates as a platform for high-resolution bioimaging. As proof of concept, we cultivate human mesenchymal stem cells on these substrates, revealing very different growth patterns compared to flat glass. Specifically, the spatial distribution of cytoskeleton proteins is vastly modified, as we demonstrate with 3D SMLM assessment.

---

Cell growth and differentiation are driven by a multiplicity of factors influencing transcriptional activity. *In vivo*, the cell phenotype is largely influenced by the extracellular matrix (ECM), which provides both a chemical and physical environment for the cells to differentiate, migrate, change phenotypes and proliferate [1, 2, 3, 4]. *In vitro*, it has thus long been a concern that cells cultivated in 2D on flat glass (or plastic) substrates often do not behave as in their native environment [5]. The differences range from differentiation of stem cells or partially differentiated cells, to growth and proliferation, to phenotypic changes and altered response to drugs, to adhesion, motility or simply cell shape [6, 3]. Still, in many cases, cultivating cells on flat substrates remains the preferential method due to its simplicity, lower cost and the availability of standardized procedures. On the contrary, *in vivo* animal experiments are often hampered by ethical concerns as well as scalability issues including high costs, extended experimentation time required to obtain the samples and reduced number of subjects. They are therefore not suited to screening large sets of drug candidates or experimental conditions. Thus, there has been an increasing need for *in vitro* advanced 3D cell culture models with the development of a comprehensive toolbox to form these models and study them [7].

Cell lines with self-assembly capabilities can be employed to create spheroids using diverse methodologies, including pellet culture, the hanging drop technique [8, 9], and the utilization of microfabricated structures such as microchannels [10] and microwell arrays [11]. Hydrogels, which have the ability to act as surrogates for the ECM, are frequently employed as matrices for encapsulating individual cells in suspension or pre-formed spheroids. These hydrogels can be tailored to incorporate specific biochemical, physical, and structural components of the ECM [12, 13, 14]. Moreover, they can be tuned to promote cell adhesion properties, which is achieved through the incorporation of arginylglycylaspartic acid (RGD) moieties to support interactions with integrins and other cell adhesion receptors [15]. Further enhancements can be achieved by patterning the hydrogel or integrating it within a microfluidic device, introducing additional mechanical cues stemming from interstitial flows or cell-induced mechanical forces [16, 17]. Still, the 3D cell culture in hydrogels can hinder visualization, particularly in high resolution imaging, therefore other methods to provide 3D cues to cells have been investigated, including the use of substrates decorated with 3D multiscale features that furthermore open new opportunities to study the impact of the environment on cell differentiation [3].

Moreover, aside from the interest for mimicking tissues [6, 18], cell line filtering and isolation [19, 18] also call for growth and differentiation control [20, 21]. Over the past years, there has been many examples of successful modulation of cell phenotype [3]. These methods broadly fall in two main categories: chemical-induced and topographical-induced modulations. The first relies on either the addition of proteins or chemicals in the culture medium, or chemical modification of the surface of the substrate to induce the desired growth behaviors, while the second is based on tailoring the geometry of the environment in which the cells are cultivated. Various properties of the substrate can be engineered to tune the growth of cells, such as the local positioning of nano- or micro-structures, characteristic sizes or spacings, anisotropy or randomness, as well as the structure geometry length scale. The type of structures range from etched substrates to fibers to micropatterned lattices [22, 1]. In the latter category, there has been a number of studies of cell culture on pillars, grooves or pits [23, 24, 25]. In this work, we will focus on more complex structures consisting of amorphous SiO_2_ 3D pyramids or fractal octahedra, which have been previously reported [26], but not yet used for high-resolution self-referenced bioimaging purposes. These multiscale inorganic substrates, which we will refer to as ‘microstructured fractal substrates’, have been shown to induce a wide variety of behaviors from a given initial set of cell line and seeding conditions [18, 20, 21]. The size, spacing and lattice type can be varied (**Fig. 1a**), and it was observed that the level of detail of the fractal structures (i.e. the number of fractal iterations, see **Fig. 1a**) had a dramatic impact on the fate of the cells both in the frameworks of differentiation of multipotent cells [20], and differentiated cells growth patterns [18]. In particular, depending on the number of iterations, cells were found to grow in either monolayers or form 3D spheroids reliably. Interestingly, using the topography of the substrate as physical cues to drive cell differentiation and proliferation readily offers promising perspectives for drug screening, cell line purification, or even the potential for electrochemical sensing to capture additional molecular cues during cell growth [18, 20, 21, 27]. However, understanding the underlying mechanisms driving the cell phenotype for a given substrate geometry requires a more thorough assessment of cell material interactions, including the distribution of proteins involved in cell adhesion (e.g., focal adhesion point) and cell morphology (cytoskeleton) at the molecular scale, for which super-resolution imaging is well suited. This, in turn, calls for the ability to optically image and colocalize the proteins of interest in 3D at high spatial resolution without the need for tedious optical calibration and alignment protocols.

**Figure 1:**
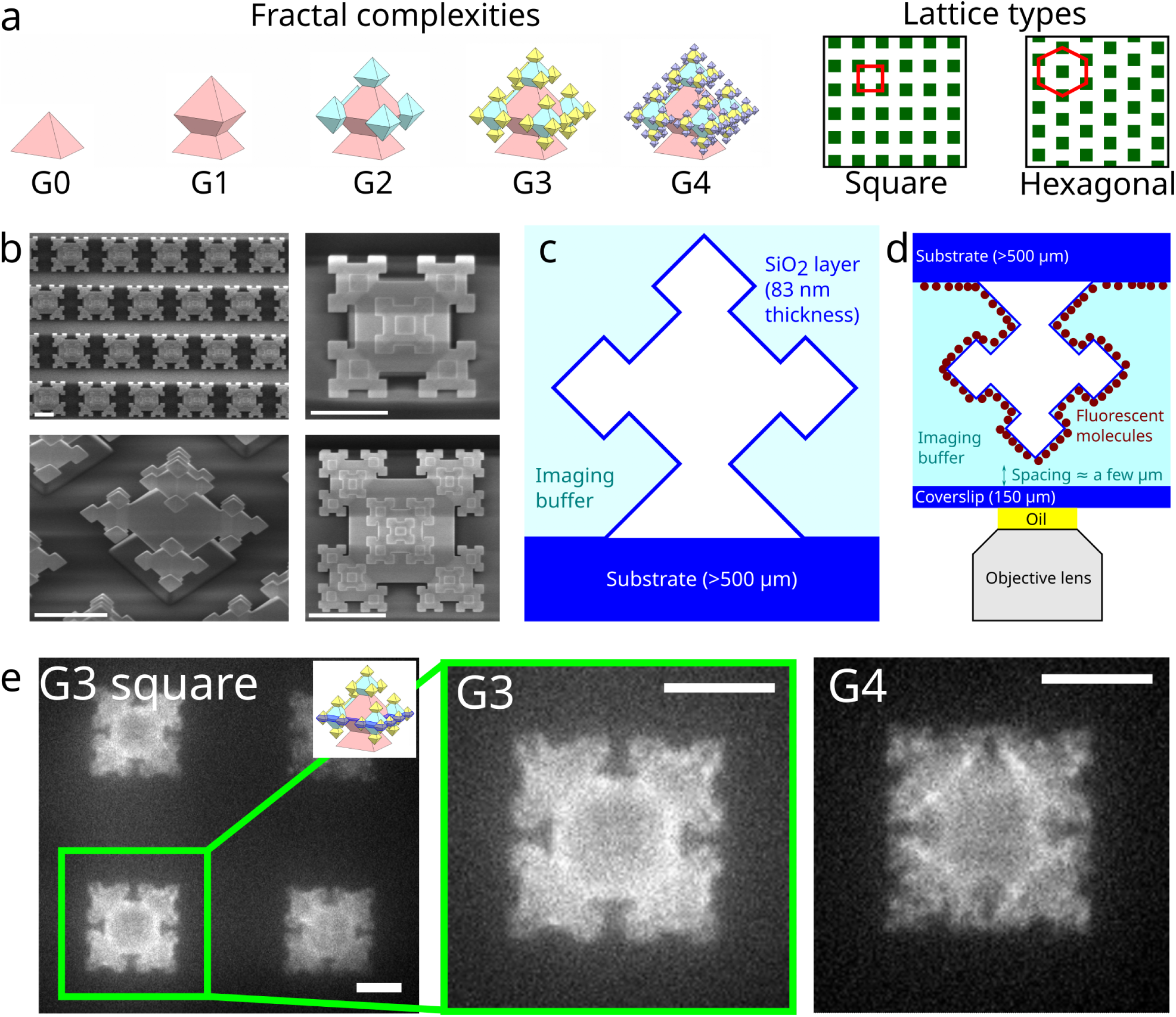
Geometry of the microstructured fractal substrates. **a** Schematic of the different fractal generations (G0–G4) and lattice types (square or hexagonal) employed in this study. **b** SEM images of G3 and G4 fractal structures on a hexagonal lattice: low (top left) and high (top right) magnification top view images of a G3 sample, (bottom left) high magnification oblique view of a G3 sample, (bottom right) high magnification top view image of a G4 sample. **c** Schematic representation of the geometry of one fractal structure (side view, slice in the middle plane). The distribution of the imaging medium is also shown. **d** Sample fluorescent labelling and mounting geometry. **e** Widefield microscopy diffraction-limited images obtained from fluorescently labelled square lattice G3 (left: large field of view, center: zoom on one fractal structure) and G4 (right: zoom on one fractal structure) samples imaged in the middle axial plane (the position of the imaging range is represented in the inset). Note that the smallest (G4) features can barely be resolved. Scale bars: 5 µm.

Here, we show that through changes to the manufacturing protocol of the 3D multiscale structures, we can achieve achieve the full potential of Single-Molecule Localization Microscopy (SMLM) to study cells cultivated on the microstructured fractal substrates. We also demonstrate how the known 3D shapes and sizes of the fractal structures and their multiscale nature can be used to both perform the required calibrations and assess the imaging resolution without the need for additional external calibration standards like fluorescent nanoparticles. Finally, as a bioimaging proof of concept, we reveal cell shape and adhesion modifications of human Mesenchymal Stem Cells (hMSCs) cells grown on microstructured fractal substrates compared to cells grown on flat glass coverslips by imaging the 3D distribution of proteins of the cytoskeleton (actin) at the sub-diffraction level. In the near future, 3D multiscale fractal structures can be used as a self-referenced tool in fluorescence microscopy, and to investigate biological signals and cell phenotype at high resolution.

## Results

### Geometry of the microstructured fractal substrates

We used SiO_2_ micropatterned substrates fabricated following the process described in [26, 28]. In short, the negative of the pyramids is formed in crystalline silicon by masked anisotropic etching. The negative of the fractal structures is produced by a series of repeated steps starting from these pyramidal pits: the sides of the hole and top surface are oxidized to create a SiO_2_ protective layer. Next the concave corners of the SiO_2_ mask are opened selectively. These openings are the starting point for the anisotropic etching to create octahedron-shaped holes. This process can be iterated to produce a series of octahedra of decreasing sizes. Eventually, oxidation, anodic bonding to a glass substrate and etching of the remaining crystalline Si wafer turns the negative mould in the convex 3D structures depicted in **Fig. 1a**. The iterative nature of the manufacturing process offers the possibility to create structures of increasing detail level in a fractal manner. The number of steps performed to etch octahedra is called the *generation* (abbreviated as *G*) of the structures. Therefore, G0 corresponds to a pyramid, while G1 adds an octahedron at the tip of the pyramid, and G2 adds one octahedron at each tip of the G1 octahedron, etc. (**Fig. 1a**). Structures up to G4 can be manufactured. Aside from the number of iterations, the type of the 2D lattice over which the fractal structures are organized can be chosen arbitrarily, and we used either hexagonal or square lattices here. Finally, the spacing between the fractal structures can be chosen, as well as their sizes. Fractal shapes are intrinsically multiscale objects, with spatial dimensions covering over a wide range of sizes. Here, the spatial period of the lattice is typically around 10 to 20 µm for the hexagonal and square lattices respectively, while the sides of the largest (G0–G1) and finest (G4) features are around 5 µm and 500 nm respectively (see **Fig. 1b**). We also investigated different sizes to validate their application as 3D multiscale self-referenced structures for SMLM.

The pyramids/octahedra are composed of a SiO_2_ layer delineating a hollow core whereas the space available to the cells and the imaging buffer is mainly water-based (see **Fig. 1c**). The resulting refractive index mismatch between the interior and exterior causes a significant amount of optical aberrations as the light travels through the material. While this has not prevented 2D diffraction-limited imaging in confocal microscopy or white light transmission [18, 20], it has led to detrimental artefacts in 3D imaging (see **Fig. 3c1** in [18]). Furthermore, super-resolution microscopy methods such as SMLM are known to be sensitive to optical aberrations that lead to a decrease of the fluorescence light intensity and a deformation of the Point Spread Function (PSF), i.e. the image of the single molecules. This often results in a deterioration of the localization precision [29, 30], and can furthermore induce axial biases in 3D SMLM [31, 32, 33, 34, 35, 36]

### Diffraction-limited 2D optical imaging of the microstructured fractal substrates

To eventually demonstrate super-resolution fluorescence imaging of specific proteins in cells grown on these substrates, the first step was to image the bare microstructured fractal substrates to determine their exact 3D sizes and verify the consistency with the manufacturing specifications. After imaging them using scanning electron microscopy (SEM) as in **Fig. 1b** to provide a reference for comparison, we performed a labelling of the glass surface using N-Hydroxysuccinimide (NHS) ester chemistry (see **Methods**) to attach photoactivatable fluorescent molecules (Abberior Cage 635). We then set out to image the labelled microstructured fractal substrates using wide-field fluorescence microscopy. However, optical microscopy imposes some constraints in terms of imaging conditions. Indeed, inverted microscopes are usually designed to image biological samples cultivated on standard 170- or 150-µm thick glass coverslips in reflection configuration. In many cases, this is not compatible with stringent fabrication requirements, which are often incompatible with such fragile substrates. Micropatterned substrates are more commonly designed on 500 µm thick wafers.

We therefore imaged the samples in a ‘flipped down’ configuration (see **Methods**, as well as **Fig. 1d**), i.e. with the pyramids/octahedra facing down and placed on a coverslip with a 150-µm nominal thickness. Note that, as a convention, the results will still be displayed facing up (i.e. as in **Fig. 1c**) rather than down in this manuscript to facilitate the comparison with SEM images. To keep the acquisition conditions as close as possible to those that we will use for cell imaging, we used Phosphate Buffered Saline (PBS) as an imaging medium. In such a configuration, the fractal structures and the bottom 150-µm coverslip are separated by a spacing of one to several µm (possibly due to the slight flatness nonidealities of the coverslip), thus preventing potential damage resulting from direct contact.

**Fig. 1e** presents images of the G3 and G4 functionalized microstructured fractal substrates obtained in wide-field microscopy (see **Methods**). Due to the limited depth of field (approximately ±500 nm relative to the focal plane), these images correspond to cross-sections of the 3D microstructures in the *xy* plane, and we imaged the middle plane, i.e. the plane that maximizes the apparent *xy* size of the cross-section. The images allow distinguishing the lattice as well as the large features (G1, G2 and G3 octahedra), however, the G4 octahedra are close enough in size (∼450 nm) to the diffraction limit (∼300 nm) that they are challenging to distinguish, calling for a method with higher spatial resolution. Furthermore, the imaging only yields the projection in 2D of the molecules located within the depth of field (∼1 µm), which highlights the need for 3D imaging.

### Super-resolved 2D optical imaging of the microstructured fractal substrates

We first modified the acquisition conditions to obtain sub-diffraction spatial information using SMLM [37, 38, 39]. By increasing the excitation power and carefully tuning the photoactivation power (see **Methods**), we achieved a blinking regime characterized by the sparsity of active fluorescent molecules in each frame of the recorded video. Using a homemade single-molecule localization code (see **Methods**), we were able to pinpoint the position of each molecule labelling the substrate with a precision in the 10-nm range. This information was used to reconstruct super-resolved pointillistic images of the labeled structures. They revealed much finer details compared to the diffraction-limited images obtained by wide-field fluorescence microscopy, while retaining the ability to faithfully report features in the range of µm or tens of µm, as shown in **Fig. 2a–b**. The SMLM cross-section image of the square lattice G1 labelled sample reveals a lattice unit length of 20.36±0.25 µm in *x*, and 20.30±0.31 µm in *y*. The dimensions are in good agreement with those measured in SEM from the same sample, which were found to be 20.66±0.26 µm and 20.26±0.32 µm in *x* and *y* respectively. As a convention, we display the results as the mean ± twice the standard deviation of the distribution of the measured values. 2D SMLM and SEM distance measurements are summarized in **Supplementary Note 1**, which also displays the sizes of the statistical sets. We point out that, interestingly, this functionalized microstructured fractal substrate could offer a viable alternative for the calibration of fluorescence microscopes, which are usually calibrated using targets of known sizes used in white light transmission, potentially inducing biases due chromatic aberrations.

**Figure 2:**
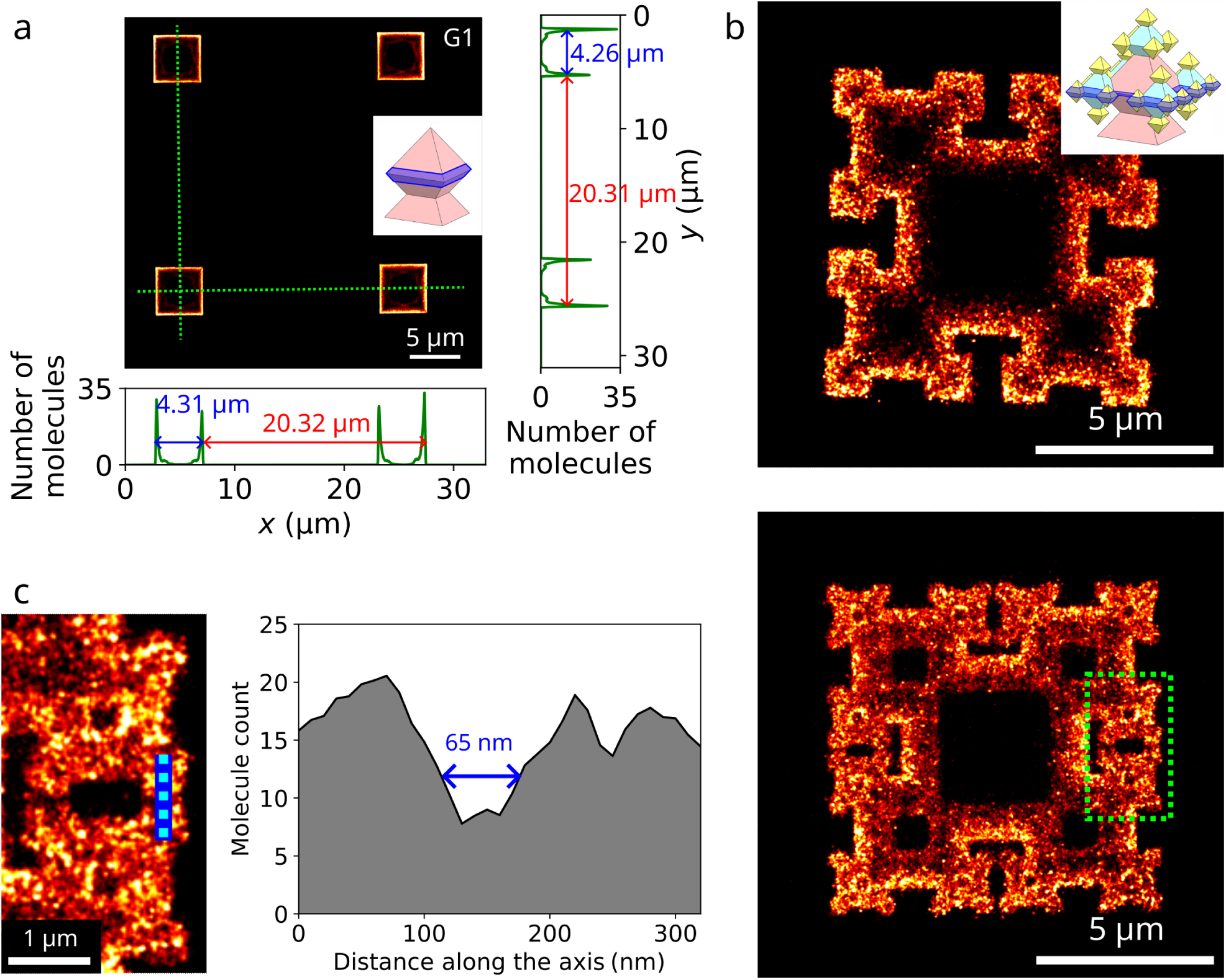
2D SMLM imaging of the microstructured fractal substrates labelled with fluorescent molecules. **a** 2D SMLM image of a square lattice G1 sample labelled with fluorophores and imaged in the middle axial plane (as depicted in the inset). The field of view contains four fractal structures. The color encodes the density of molecules. *x* and *y* molecule density profiles are plotted along the green dotted lines, which allow the measurement of the lattice period and the size of the G1 octahedron. **b** 2D SMLM molecule density images of square lattice G3 (top) and G4 (bottom) microstructured fractal substrates labelled with fluorophores and imaged in their respective middle planes (see inset). Each image shows a zoom on one single fractal structure, which allows to clearly resolve the G0–G4 octahedra. **c** Zoom on the area indicated with a dotted rectangle in **b** and molecule density profile along the cyan dotted line showing an illustration of the resolution power of SMLM from the gap between adjacent G4 octahedra. Scale bars: 5 µm (**a–b**), 1 µm (**c**)

The edge length of the G1 octahedron imaged in its middle plane (see **Fig. 2a**) was found to be 4.34±0.24 µm and 4.32±0.12 µm in *x* and *y* respectively in SMLM, compared to 4.39±0.12 µm and 4.41±0.14 µm in SEM, again in good agreement. To further verify the reliability of our SMLM strategy, we imaged functionalized microstructured fractal substrates of higher generations (see **Fig. 2b**). We found the G3 features to be 1327±72 nm and 1295±74 nm in *x* and *y*, compared to 1330±28 nm and 1299±29 nm in SEM. Finally, the G4 features were measured 453±26 nm and 449±29 nm in *x* and *y*, compared to 457±22 nm and 448±20 nm in SEM.

Finally, to demonstrate the possibility to image down to the sub-100-nm scale, we investigated the gap between adjacent G4 features in the middle plane (see **Fig. 2c**). While this gap may vary in size from one microstructure to the other or between several pairs of G4 octahedra among the same microstructure due to fabrication nonidealities, it is consistently below 200 nm, i.e. in the sub-diffraction domain. 2D SMLM allowed to resolve all the gaps investigated and the spacing value was measured down to 65 nm, which validates the resolution of the imaging technique in spite of the challenging acquisition conditions. The measured gap distances are once more consistent with SEM results (see **Supplementary Note 1**). This demonstrates that the fluorescently-labelled microstructured fractal substrates can be used as calibration tools to allow multiscale resolution measurements, particularly in the framework of high resolution bioimaging of cells cultivated on such substrates, which will be investigated later in this manuscript as a proof of concept.

We conclude from these characterizations that the SMLM approach is able to accurately capture submicron features, down to the scale of a few tens of nm, while retaining the capacity of wide-field imaging to render larger features in the range of tens of microns. We further conclude that SMLM can be used as an affordable substitute for SEM to characterize glass samples over a range of lengths spanning a hundred nanometers to several tens of microns. Importantly, SMLM does not rely on destructive specimen preparation and is not limited to surface imaging. Moreover, it can be used for specific observation, i.e. different chemical species can be labelled with different fluorescent molecules and subsequently distinguished, which is a challenging task in SEM. Finally, unlike SEM, SMLM is a priori compatible with imaging of living cells and biochemically functionalized surfaces. The typical time required to obtain an SMLM image is around 15 min. In both SEM and SMLM, the resolution is comparable. On the one hand, SMLM can achieve a resolution around 10 nm, limited by the localization precision and the size of the label. On the other hand, SEM has similar resolution limitations due to glass charging.

### 3D single-molecule imaging of the microstructured fractal substrates

Another major advantage of SMLM is its ability to provide information about the position of the molecules along the optical axis. Several approaches exist to return the axial position, the most common being the measurement of the shape of the PSF [40, 41, 42]. Other strategies include interferometric measurements [43, 44, 45], molecule triangulation [46, 47], interface effects [48, 49, 50] or combinations of several of these approaches [51].

We set out to obtain a 3D rendering of the fractal microstructures through SMLM. We employed the widely used astigmatic imaging approach [40] (see **Methods** as well as **Supplementary Fig. 1** for a schematic of the optical setup). It has the advantages of implementation and processing simplicity, versatility and extensive use in the SMLM community, making it a standard method. The downsides are that it does not provide as precise axial localization as other approaches, especially far from the focus plane [42, 43, 52, 45], and that it requires a subtle calibration step. Indeed, due to the index mismatch between the substrate (*n*_glass_ ≈ 1.5) and the aqueous imaging medium (*n*_water_ ≈ 1.33), a depth-dependent amount of spherical aberration adds to the defocus and the static optical aberrations induced by the detection system. As a consequence, the shape of the PSF is dependent on both the position of the emitter relative to the focus and on the position of the focus relative to the substrate. While these nonidealities can be challenging in many cases, the known 3D geometry of the fractal structures constitutes a valuable tool to tackle them.

We first performed a calibration of the astigmatism using beads deposited on a flat glass coverslip and flipped down as depicted in **Fig. 1d** (see **Methods**). The piezoelectric stage was scanned axially at known positions. This provided the correspondence between the shape of the PSF and the depth relative to the focal plane. Then we proceeded with the 3D imaging of the labelled microstructured fractal substrates. Unlike more refined PSF shape measurement approaches [53], astigmatism-based 3D SMLM, however, does not have the capacity to image the full height of the fractal structures (between 3 and 8 µm, depending on the chosen geometry) at a given focus plane, since the axial range of imaging is limited by the shallow depth of field. More precisely, only molecules located within approximately ±500 nm relative to the focus plane can be localized. We first performed imaging of only a cross-section of the microstructured fractal substrates at a given height. However, contrary to the 2D SMLM images presented in **Fig. 2**, we now have access to the axial positions of the molecules imaged in the depth of field. **Fig. 3a** shows the 3D images of the same field of view of a functionalized hexagonal lattice G3 sample slightly above the middle plane, slightly below the middle plane and slightly above the flat bottom substrate respectively (the axial position is color-coded). It can be seen that 3D SMLM retains the ability to capture the *xy* features with a good precision. In addition, the 3D positions can be used to determine the local slope. Above the middle plane, the positions range from lower values (red) at the edges to higher values (blue) towards the center of the squares. Below the middle plane, on the contrary, the fractal structure indicated with a cyan dotted square shows the opposite behavior, with lower heights towards the center compared to the edges. Finally, close to the flat bottom substrate, the same behavior is observed than above the middle plane, i.e. the height increases towards the center. The flatness of the bottom substrate is also evidenced. All these observations are consistent with the expected shape (see insets).

**Figure 3:**
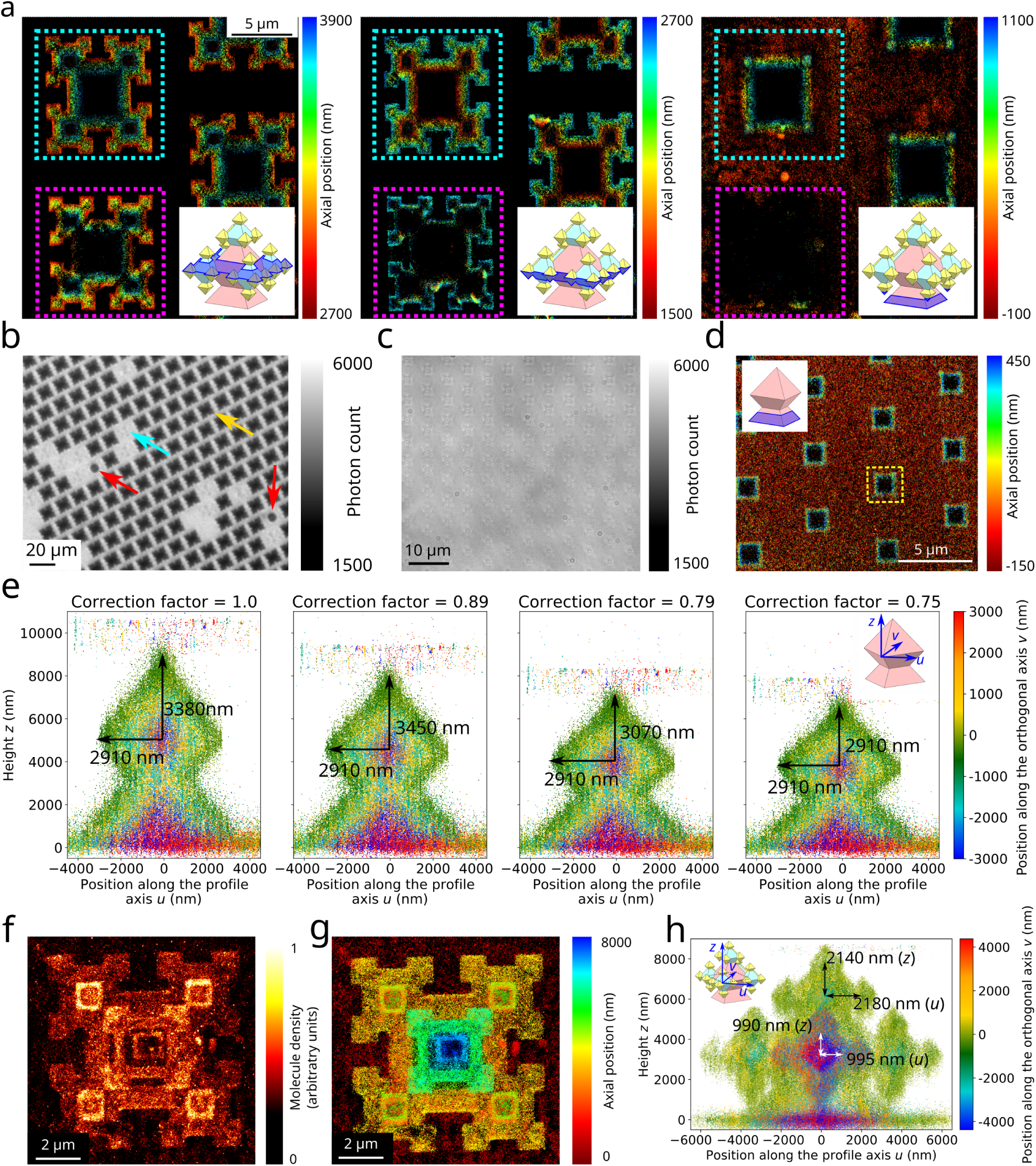
3D SMLM imaging of the microstructured fractal substrates labelled with fluorescent molecules. **a** 3D SMLM of a hexagonal lattice G3 microstructured fractal substrate functionalized with fluorescent molecules and imaged at three different focus plane positions (the corresponding capture ranges are indicated by the insets): slightly above the middle plane (left), slightly below the middle plane (center) and slightly above the flat bottom substrate (right). The height of each molecule is color-coded. The fractal structures indicated with cyan and magenta dotted squares can be imaged over the whole volume or only over the upper half, respectively. **b** White light transmission microscopy images of a hexagonal lattice G3 sample. Note the three different populations of fractal structures: those filled with air (yellow arrow), those completely filled with imaging buffer (cyan arrow) and those partially filled with imaging buffer (red arrows). **c** White light transmission microscopy images of the variant of the hexagonal lattice G3 microstructured fractal substrate featuring perforated tips. Note that the microstructures are almost invisible. **d** 3D SMLM image of the hexagonal lattice G3 sample with perforated tips labelled with fluorescent molecules. The imaging plane is set slightly above the flat bottom substrate, as depicted in the inset. The yellow dotted square indicates the approximate shape of a fractal structure in the middle plane, which highlights that there is no shadowing effect occurring for the molecules below the middle plane. Note that the perforated fractals were manufactured with different sizes than the non-perforated. **e** 3D SMLM side view of a G1 microstructure labelled with fluorescent molecules, filled with imaging buffer, and imaged by continuously scanning in *z*. The molecules are projected on *uz*, in the plane of the diagonal (see the inset on the right) and the position along the perpendicular direction *v* is color-coded. Several commonly used values are assumed for the focal shift correction factor (1, 0.89 and 0.79), resulting in a more or less visible stretching of the octahedron in the axial direction. The value 0.75 provides an exact matching of the lateral and axial diagonals, i.e. a slope of 45°. **f–g** 2D (**f**) and 3D (**g**) SMLM images of the same G3 fractal structure labelled with fluorescent molecules and filled with imaging buffer. The full extent of the microstructure was imaged by continuously scanning the focal plane along the axial dimension. The color encodes the density of molecules (**f**) or the axial position of each molecule (**g**). **h** 3D SMLM side view of the G3 microstructure presented in **h** for a focal shift value of 0.75. The molecules are projected in *uz*, in the diagonal plane (see the inset). The position along the perpendicular axis *v* is color-coded. The lateral and axial sizes of the G2 and G3 features are displayed. Scale bars: 5 µm (**a,d**), 20 µm (**b**), 10 µm (**c**), 2 µm (**f,g**).

Interestingly, the fractal structure indicated with a magenta dotted square in **Fig. 3a** is as visible as the first above the middle plane but seems to disappear slightly below the middle plane, and is completely invisible close to the flat bottom substrate. Besides, it can be seen that the molecules attached to the flat bottom substrate do not appear whenever they are located in the dotted square, i.e. directly below the shape that the microstructure has in the middle plane. To understand the origin of this shadowing effect, we investigated the blinking video recorded, as shown in **Supplementary Video 1**, which revealed that the fractal structure indicated with a cyan dotted square displays PSFs suitable for 3D localization regardless of the imaging plane. The fractal structure indicated with a magenta dotted square displayed very similar PSF shapes slightly above the middle plane, but distorted and widened shapes slightly below the middle plane. Finally, slightly above the flat bottom substrate, almost no PSF could be observed. This suggests noticeable optical aberrations, which implies that the fluorescence light is affected by a refractive index mismatch along its propagation from the molecule to the objective.

We hypothesized that the refractive index mismatch may be due to the fact that the fractal structures are composed of a thin SiO_2_ layer enclosing a hollow volume. Assuming that the fractal structures are perfectly closed, that space (depicted in **Fig. 1c–d**) should be filled with air in our acquisitions. However, defects in the fabrication process or mechanical pressure induced during the handling of the sample may create local holes or cracks that allow the imaging medium to flow through and fill the space inside some of the octahedra, thus effectively making the 3D structures transparent, as the thickness of the glass layer (∼83 nm) and the index mismatch (Δ*n* ≈ 0.17) are too low to induce a noticeable amount of optical aberration. If this hypothesis is valid, then the refractive index mismatch should be visible in white light transmission microscopy. We therefore prepared a hexagonal lattice G3 sample mounted in the same configuration and with the same imaging medium as previously, and we imaged it at low magnification with white light transmission microscopy. **Fig. 3b** allows to distinguish three populations of fractal structures: octahedra filled with air appear in black due to the diffraction induced by the index mismatch, while octahedra filled with imaging buffer (in gray) are barely distinguishable from the flat bottom substrate between the fractal structures because the light does not undergo significant refraction or scattering. Finally, a small subset of the octahedra appear partially filled with imaging buffer. Time-lapse imaging with white light transmission microscopy (see **Methods**) of this third population shows that they slowly become filled with imaging buffer over the course of several tens of minutes (see **Supplementary Fig. 2**). Mechanical stress (i.e. gentle squeezing) prior to immersion resulted in a rapid increase of the proportion of fractal structures being filled with imaging buffer within a few minutes only. From these results, we conclude that in **Fig. 3a**, the fractal structures indicated with cyan and magenta dotted squares were filled with imaging buffer and air respectively. It should be noted that only the light emitted by molecules located behind the fractal structures, i.e. below the middle plane, is affected by the diffraction due to the index mismatch.

While we were able to image some of the fractal octahedra due to structural defects, we point out that a number of them remain impossible to image below the middle plane. Induced mechanical stress is not a viable method as it may result in significant damage to the microstructures. We therefore chose to modify the fabrication protocol in order to etch holes at the tips of the smallest octahedra, using a technique reported in [26]. This hole opening at the tip has been possible by keeping the SiO_2_ mask as the final oxide thickness after isotropic etching and selectively opening the mask before bonding to the glass substrate and etching the remaining crystalline Si wafer, which leaves orifices at apexes. We thus manufactured a hexagonal lattice G3 microstructured fractal substrate with perforated tips (see **Supplementary Fig. 3** for the SEM characterization). Note that the sample was manufactured with smaller sizes (edge lengths of approximately 1630 nm, 790 nm and 370 nm for the G1, G2 and G3 octahedra respectively), though the fabrication process is not limited with regards to the size of the structures. White light transmission microscopy confirmed that all the octahedra are completely filled with imaging buffer, essentially making them almost invisible (see **Fig. 3c**). Accordingly, PSFs obtained in the SMLM regime after functionalization of the microstructured fractal substrates were found to be unaffected by the optical aberrations previously evidenced, as shown in **Supplementary Video 2**. This allowed us to perform SMLM imaging close to the flat bottom substrate (**Fig. 3d**), which demonstrates that 3D super-resolution can be achieved across the full axial extent of all the fractal structures on the substrate.

### 3D SMLM axial calibration

Although the possibility to image any plane in 3D SMLM paves the way for the 3D reconstruction of the whole fractal structures ultimately, it is first necessary to accurately calibrate the axial position of the imaging plane. Indeed, it is known in optical microscopy that the displacement of the focal plane is not equal to the displacement of the piezoelectric stage when scanning the sample in the axial direction. This is due to the refraction of the beam undergone between the imaging medium (*n*_water_ ≈ 1.33) and the glass coverslip (*n*_glass_ ≈ 1.5). The determination of the correction factor, called *focal shift* has been investigated by different groups in the context of fluorescence microscopy, and experimental publications [54, 36] suggest that the approaches derived from ray optics approximations resulting in values of *n*_water_/*n*_glass_ ≈ 0.89 or (*n*_water_/*n*_glass_)^2^ ≈ 0.79 used in the early days of 3D SMLM (for instance in [40]) are not valid for the high numerical aperture objectives used in SMLM. However, the experimental assessment of the focal shift is no small task as it relies on the hypothesis that the fluorescent sources are at known axial positions in the calibration sample.

Capitalizing on the knowledge of their geometry, we used the fractal structures themselves as a reference to measure the value of the focal shift. Indeed, during the manufacturing process, the wet etching step produces faces aligned with the ⟨100⟩ lattice [26]. In other words, the angle between the edges and the horizontal plane is expected to be 45°, and all the diagonals should have the same length.

In order to capture the full height (up to 8 µm) of the fractal octahedra, we collected 3D SMLM data at different focal plane positions along the optical axis by moving the piezoelectric stage on which the sample is placed. By stacking the results, we obtained an image over an extended axial range. While unusual in SMLM, this strategy was successfully applied in [34]. Rather than imaging at a few discrete values of the focal plane position and stitching them together, which has the disadvantage of yielding a strongly depth-varying axial precision, we chose to apply a smooth focal plane displacement during a long continuous acquisition by shifting the piezoelectric stage by 100-nm steps (see **Methods** for more details about the acquisition). The uncorrected results (i.e. for a focal shift factor of 1) of a full scan of a G1 microstructure are displayed as a 3D SMLM side view in **Fig. 3e left**. The octahedron displays a noticeable stretching in the axial direction, resulting in an axial diagonal seemingly larger than the lateral diagonal, which highlights the impact of the focal shift on the 3D data. Correction factors of 0.89 and 0.79 (i.e. *n*_water_/*n*_glass_ and (*n*_water_/*n*_glass_)^2^ respectively) reduced the apparent stretching without fully correcting it (see **Fig. 3e center left** and **center right**). Optimal match between the axial and lateral diagonals was achieved for a correction value of 0.75 (as depicted in **Fig. 3e right**). This experimentally determined value of the focal shift is consistent with the values determined from [54] (0.73) and [36] (0.75–0.80) for our experimental conditions.

Interestingly, the known geometry of the microstructured fractal substrates and their suitability for cell culture make them valuable tools for self-referenced measurements. Indeed, calibration steps such as the determination of the focal shift, as well as the generation of the astigmatism curve, can be performed on the same type of substrates as the fluorescence imaging of the biological samples of interest. This reduces the sources of systematic error that can arise from potential discrepancies between the calibration and imaging conditions.

Acquisitions performed on G3 structures assuming this measured value of the focal shift highlight the capacity of this technique to faithfully capture fine structures over the full extent of the 3D substrates well beyond the range of the depth of field both in the lateral and axial directions (**Fig. 3f–g**). In particular, we found the lateral diagonal to be equal to the axial diagonal for each generation of octahedra investigated, as illustrated in the 3D profiles displayed in **Fig. 3h**.

It should be noted that this quantitative characterization is easier to perform in 3D SMLM than in SEM since the former produces actual 3D results. This makes rotations of the structures over arbitrary axes easy as it only requires a modification of the display (see **Supplementary Video 3**). Similarly, slicing and projecting in arbitrary planes are straightforward post-processing steps, as illustrated in (see **Supplementary Fig. 4**). On the contrary, SEM essentially returns 2D images that are projections over planes defined during the acquisition. Lastly, imaging glass surfaces behind the first encountered is not possible in SEM, while the perforated fractal structures effectively behave as a homogeneous transparent medium in optical imaging, allowing to image molecules on different surfaces beyond the first (as in **Fig. 3a,e,h**).

### High resolution 3D bioimaging

Having validated the precision, reliability and versatility of 3D SMLM imaging on the microstructured fractal substrates of known sizes decorated with fluorescent molecules, we eventually set out to image cells cultivated on them in 3D SMLM. As mentioned previously, the microstructured fractal substrates are of particular interest in cell biology due to their capacity to modulate the cell phenotype and behavior. This was successfully used to isolate cell lines, promote tumor growth and drive cell differentiation in a fast and reliable way [18, 20]. More generally, 3D substrates open promising perspectives to produce complex *in vitro* models faithfully mimicking cell behaviors in tissues that can be subsequently used for a range of applications, particularly in the domain of drug screening and toxicity measurements. While 2D diffraction-limited imaging has been achieved on the microstructured fractal substrates with white light transmission or confocal fluorescence modalities [18, 20], super-resolution has not been demonstrated so far, and 3D diffraction-limited imaging has proved prone to imaging artifacts [18]. However, given the results that we obtained so far in 3D SMLM, as well as our capacity to produce fractal octahedra with perforated tips to adapt the optical index, high resolution 3D cell imaging has now become reachable.

Human mesenchymal stem cells have the capacity to differentiate into osteoblasts, chondrocytes, adipocytes and many other cell types, making them a versatile cell source for tissue regeneration and regenerative medicine [55]. Micro- and nanopatterned surfaces provide cues for cell motility [56], adhesion, survival, viability and lineage specification, potentially impacting the fate of hMSCs [57, 58, 59]. Understanding how topography directs especially osteogenic differentiation is a main focus in bone tissue engineering [60, 61].

We seeded hMSCs on both flat glass substrates and square lattice G3 or G4 microstructured fractal substrates and grew them in the same conditions. No differentiation (neither osteogenic nor adipogenic) was observed on any of the surfaces over two weeks (see **Methods**). Fixation was performed after 3 days. A first visual inspection with white light transmission imaging revealed that the cells grew in an isotropic manner on the flat glass substrates. More specifically, they appeared flat and many of them were stretched in seemingly random directions, reaching sizes up to 100–150 µm. On both G3 and G4 surfaces, cells were found to align with the pattern of the microstructured fractal substrate within 3 days of culture. No significant difference was observed between G3 and G4 surfaces. In both cases, the cell density seemed variable from one area to the other.

We labelled the cells with phalloidin-Alexa Fluor (AF) 647 to stain the polymerized F-actin cytoskeleton. This labelling provides information about the general shape of the cells, and can further highlight the adhesion behavior since actin is particularly visible in focal adhesions [62, 63]. Preliminary diffraction-limited inspection performed with wide-field fluorescence imaging (by flipping down the sample as previously, see **Methods**) of the samples confirmed the trends observed in white light transmission microscopy. hMSCs cultivated on flat substrates exhibited sizes often superior to the imaging field of view (above 130 µm) and formed actin projections in seemingly random directions characterized by visible fiber bundles (see **Fig. 4a–b**). Some 3D information can be obtained by acquiring axial stacks of wide-field images, although the axial resolution is limited around 0.6 µm. We thus observed that in dense regions, the cells tended to grow on top of each other, effectively creating a multi-layer accumulation of relatively flat cells (see **Supplementary Video 4**). When cultivated on microstructured fractal substrates (whether G3 or G4), on the other hand, the cells seemed to exhibit various behaviors with two main trends: at lower densities, they tended to grow in a very anisotropic way, following the grooves between the fractal structures, either along the axes of the lattice or in diagonal (see **Fig. 4c–d** and **Supplementary Video 5 left**), often wrapping around the fractal structures (see **Supplementary Video 5 left**). They were also frequently found to grow preferentially on the fractal structures and following the axes of the lattice, sometimes using them as anchor points to remain suspended above the flat bottom (see **Supplementary Video 5 center**). In all these cases, they were found to be elongated, with lengths above 130 µm, but also very narrow along the perpendicular axis, with widths typically around the size of the spacing between two fractal structures (∼12 µm). At higher densities they favored growing as a layer resting on top of the microstructures (see **Supplementary Video 5 right** and **Fig. 4f**), effectively restoring some isotropy in their directionality. Qualitatively, no significant difference was observed between G3 and G4 microstructured fractal substrates. Interestingly, at the edge of the substrate, the interface between the area decorated with microstructures and the flat area seemed to hamper cell spreading, as the hMSCs mostly remained on the microstructures without settling on the flat part of the substrate (see **Fig. 4e–f**).

**Figure 4:**
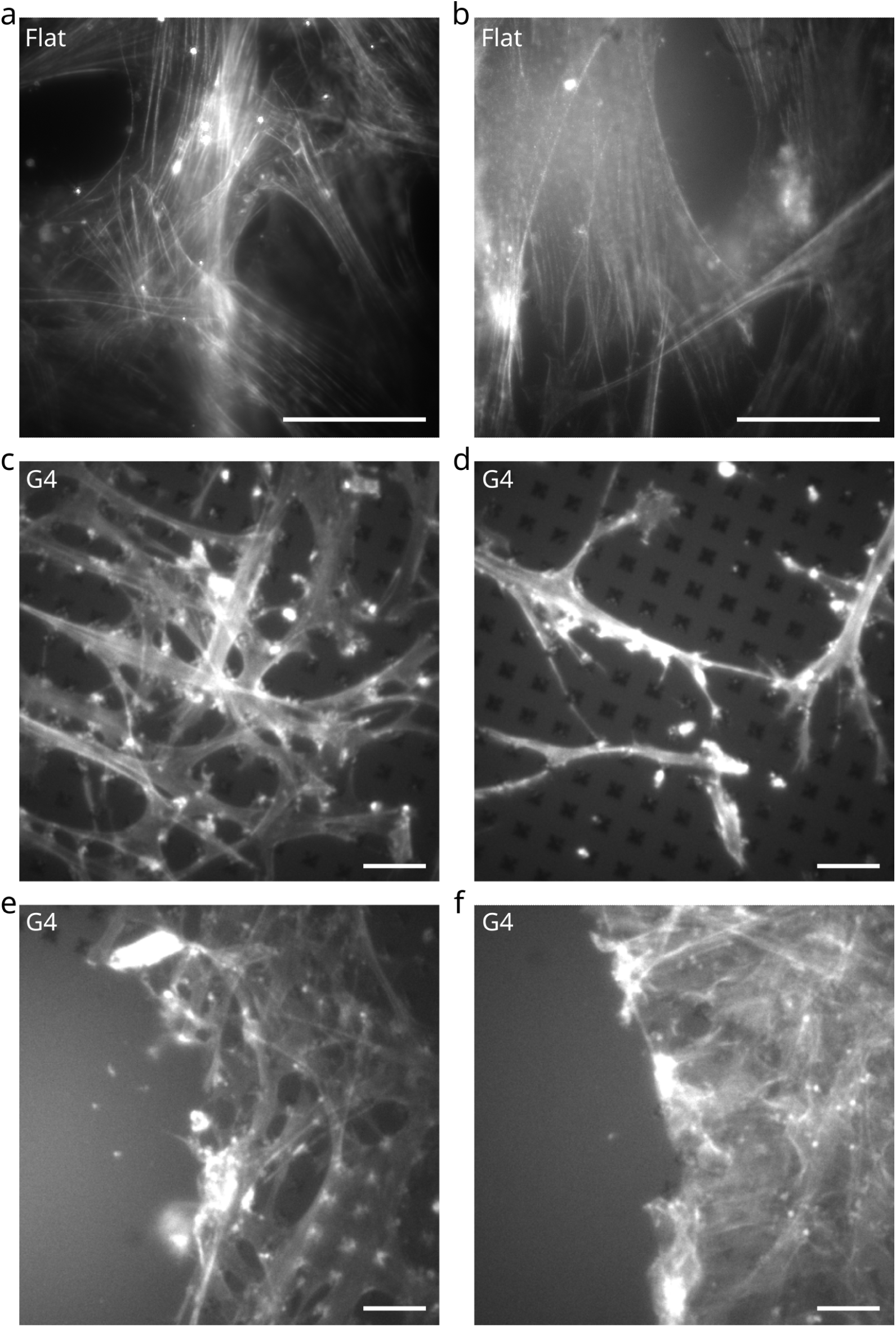
Wide field fluorescence microscopy images of hMSCs. The hMSCs were fixed after 3 days and the F-actin cytoskeleton was labelled with phalloidin-Alexa Fluor 647. The images are 2D and diffraction-limited but show the general shapes of the cells on different substrates over large fields of view. **a–b** hMSCs cultivated on a flat glass substrate, exhibiting isotropic shapes. The bright hotspots are fluorescent beads (not used for the wide field fluorescence microscopy imaging but only for SMLM imaging). **c–f** hMSCs cultivated on a square lattice G4 microstructured fractal substrate. The axes of the lattice are oblique, visible thanks to the periodic dark squares. Note that the cells seem to grow along the axes of the lattice and preferentially on the fractal structures, avoiding the flat part of the substrate (**e–f**). Note also that when the cells are dense (**f**), they tend to form one or several layers above the microstructures, which makes them grow more isotropically. Scale bars: 40 µm.

SMLM acquisitions were performed in conditions mostly similar to previously when imaging the functionalized microstructured fractal substrates (see **Methods**). In particular, the sample was mounted in flipped down configuration, and a high excitation laser power was used to induce blinking of the molecules. A notable difference compared to previous acquisitions is the chemical composition of the imaging buffer, which contains oxygen-scavenging reagents to prevent photobleaching [64, 65]. We used the same PSF-shaping-based 3D SMLM detection modality relying on the use of astigmatism, and imaging was performed at a fixed focus plane position.

The super-resolved results revealed details that are not visible on the diffraction-limited wide-field fluorescence images. hMSCs cultivated on flat glass substrates exhibited well-defined stress fibers, as shown with magenta arrows in in **Fig. 5a**. While those are also visible in wide-field microscopy, finer filaments and other features (indicated with cyan arrows) such as pseudopodia require superresolution to be identified. The 3D image (**Fig. 5b**) revealed a very flat shape of this part of the cell, characterized by a distribution of the molecules over a mere 400 nm above the coverslip (see **Fig. 5c**). hMSCs cultivated on G3 microstructured fractal substrates, on the other hand, displayed a very different actin distribution. Major stress fibers were not observed (as shown in **Fig. 5d–g**), and seemed to be replaced in some cases with dense bundles of smaller fibers (see **Fig. 5f–g**), below the diffraction limit. Interestingly, cells that wrap around the fractal octahedra were found to exhibit some accumulation of F-actin near the contact surfaces, as well as some bulge-like structures (indicated by the green arrows on **Fig. 5d,f**), although no characteristic size or shape consistently emerged. This is reminiscent of [25], where the labelling of the membrane of U2OS (human osteosarcoma) cells revealed similar shapes close to nanopillars. Looking at the 3D data (**Fig. 5e,g**), the cells further exhibited a much less flat shape compared to those cultivated on flat substrates, as quantified in **Fig. 5c**. This is also manifest on the 3D SMLM images at the contact point with the fractal octahedra, where the cells follow the shape of the 3D substrate features (the height increases towards the center of the octahedra in **Fig. 5e**, which is consistent with the slope when imaging slightly above the middle plane). The volume of the cell is visible in the groove between the microstructures too (see **Fig. 5g**), where fibers seemed to frequently stretch from top to bottom or the contrary (red arrows) and some quasi-cylindrical actin projections were observed (blue arrow). Overall, the observed 3D distributions of actin for hMSCs on microstructured fractal substrates (**Fig. 5c**) were mostly limited by the depth of field (∼1000 nm) of the 3D SMLM imaging at a fixed focal plane, while for hMSCs on flat glass substrates, they were rather limited by the flat shape of the cells (∼400 nm).

**Figure 5:**
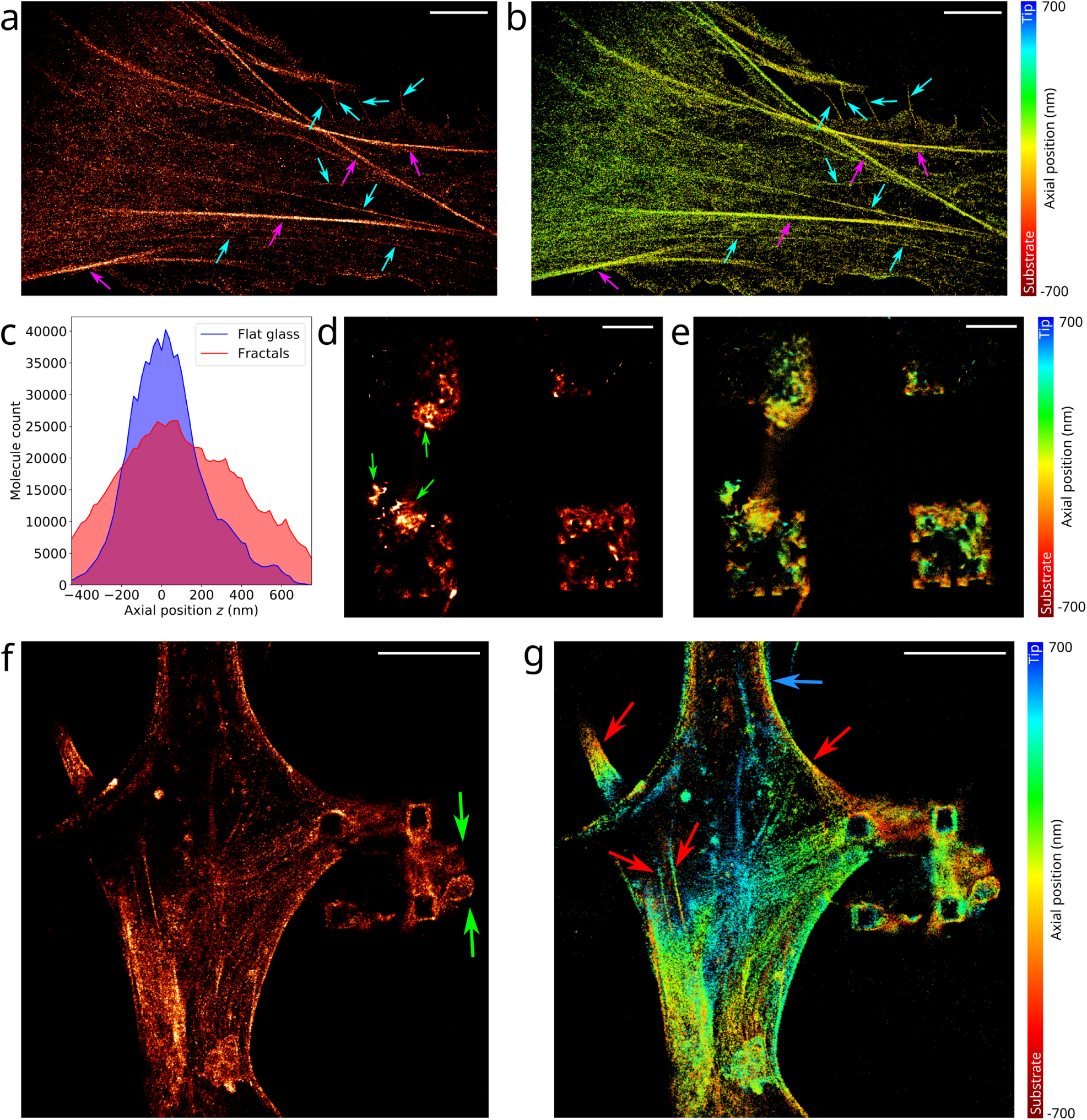
3D SMLM analysis of hMSCs fixed after 3 days and labelled with phalloidin-Alexa Fluor 647. **a–b** 2D and 3D images of cells grown on flat glass substrates: **a** displays the density of molecules, and **b** the axial position of the molecules relative to the focal plane. The magenta and cyan arrows represent structures that are already visible on the diffraction-limited images, and structures that are only visible on the super-resolved images respectively. **c** Histograms of the axial positions of the molecules in on flat glass substrates (**b**) and G3 microstructured fractal substrates (**g**). **d–g** 2D and 3D images of cells grown on a square lattice G3 microstructured fractal substrate and imaged above the middle plane. Two fields of view are shown (**d–e** and **f–g**), and the images represent the density of molecules (**d,f**) and the axial position of the molecules (**e,g**). The field of view in **d–e** contains a single cell, located mostly above the imaged plane, which forms actin projections downwards, wrapping around the fractals in the imaging plane (note that the two fractals on the top are only partially covered by the cell). The green arrows indicate accumulation of F-actin close to the surface of the octahedra as well as bulge-like structures visible only on cells cultivated on microstructured fractal substrates. Red arrows show fibers that are stretched from the top to the bottom. Finally, the blue arrow shows the almost cylindrical shape of the cytoskeleton in the grooves between the fractal structures. All axial positions are given relative to the focal plane. Scale bars: 5 µm.

Quantifying the modifications of size, shape and actin density induced by the culture on microstructured fractal substrates compared to flat glass substrates would require a more thorough and systematic analysis, but we demonstrated here the feasibility of SMLM imaging on cells cultivated on microstructured fractal substrates. It enables reliable quantitative protein distribution assessment in 3D at the super-resolution level, readily revealing features that would not be accessible in wide-field or confocal imaging.

## Discussion

We have demonstrated how 3D microstructured fractal substrates made of glass octahedra of decreasing sizes can be imaged in SMLM, particularly for 3D studies. The shape of the substrate itself can be revealed using a chemical functionalization to attach fluorescent molecules to the glass. This allows a precise and faithful characterization of the dimensions of the octahedra, therefore unlocking an alternative way of screening the substrates besides SEM. As 3D SMLM essentially returns actual 3D information in the sense that the axial and lateral position of each molecule is recorded, it is more suitable than SEM to 3D analyses relying on axial slicing or scanning, or projections along arbitrary planes.

Following an inverse approach, we also proposed the use of these microstructured fractal substrates as a calibration tool in SMLM. This is especially relevant as they exhibit three interesting features: they are 3D objects with intrinsic isotropy (i.e. the axial and lateral distances are the same), they have known sizes and geometries, and their characteristic lengths range over two orders of magnitude, from a few tens of µm to a hundred nm. We have demonstrated that they can be used to determine the value of the focal shift linking the mechanical displacement of the sample to the actual position of the focus plane in the context of axial sample positioning. We finally suggest that these samples can be used as a means to perform axial calibrations at various depths in the sample, as the spherical aberration is known to cause a depth-varying deformation of the PSFs adding to the engineered shape and potentially inducing noticeable axial localization biases in 3D SMLM if unaccounted for.

These calibration steps are relevant since they are performed in the same conditions as, and thus set the stage for, 3D high resolution of cells cultivated on the microstructured fractal substrates. The relevance of 3D substrates for cell culture has led to many publications focusing on cell proliferation and differentiation, as well as mechanobiology. Our 3D microstructured fractal substrates in particular have been previously imaged in diffraction-limited optical microscopy using white light transmission and confocal microscopy to study cell differentiation, tumour isolation and cell line purification [18, 20] at the cellular or sub-cellular scales, down to 250 nm. Here, we have demonstrated that optical imaging of cells grown on these substrates can be extended to super-resolution fluorescence imaging, in particular using 3D Single-Molecule Localization Microscopy to allow the understanding of biological matter at the macromolecular scale. While we did not perform a systematic biological study, we observed that the microstructured fractal substrates significantly modulate the growth of hMSCs, altering their shapes and sizes as well as their 3D distribution when compared to flat glass substrates. 3D SMLM revealed the reorganization of the F-actin cytoskeleton, characterized by the disappearance of large stress fibers and the accumulation of high quantities of polymerized actin around the fractal octahedra into structures lacking characteristic shapes and sizes.

Our findings open a number of interesting perspectives, either for sample characterization, SMLM calibration or biological studies. Other geometries of 3D substrates could be investigated. Notably, hollow structures made of transparent material such as SiO_2_ can be imaged in volume provided the imaging medium fills the core. It should also be noted that SMLM could be used as a means to discriminate different chemical species through the functionalization of different species with different colors of dyes.

Although we were able to demonstrate 3D SMLM on the microstructured fractal substrates, we still envision several ways to improve its performance. In particular, the index mismatch between the imaging medium and the immersion oil causes some spherical aberration that results in a loss of photons and a broadening of the PSF, and consequently a deterioration of the localization precision. Changing the objective for a silicone-immersion lens could mitigate this effect and improve SMLM results, as mentioned in [25]. Alternatively, while we have limited this proof of principle to PSF-shaping-based SMLM, other super-resolved imaging modalities can also be expected to perform well. This would be the case of techniques based on a structured excitation beam like Minflux [47] or ModLoc [45]. These techniques have provided some of the most precise and/or accurate 3D superresolution results so far.

On a different note, the multiscale and 3D aspects of the microstructured fractal substrates studied here make them promising candidates for calibration studies in SMLM or other super-resolved fluorescence imaging techniques. As we mentioned in the manuscript, they can serve as a resolution measurement tool, or an alternative to white light transmission targets to calibrate lateral distances and optical system magnifications in the specific context of fluorescence microscopy. The ability to measure the focal shift thanks to the labelled fractal microstructures is of particular interest as a recent publication suggests that the value of the focal shift may be depth-dependent [36]. The microstructured fractal substrates can furthermore be useful to generate the axial localization calibration curves at different depth values for 3D SMLM approaches relying on PSF engineering, which is already often a challenging task to perform at a single depth value [31, 32, 33, 34, 35, 36]. Finally, owing to the periodicity of the lattice, they can be used to assess field-varying aberrations and their impact on 3D SMLM, or ultimately to generate *xy*-dependent axial calibration curves, similar to [66]. Although each of these calibration steps has been studied separately, they have not been addressed simultaneously due to the lack of a suitable sample, and we believe that the microstructured fractal substrates are versatile enough to make such a goal possible.

Lastly, these multiscale substrates hold out numerous promises for cell biology. Previous studies have demonstrated their potential for cell isolation, spheroid culture or cell differentiation driving [18, 20]. The proof of principle of 3D and super-resolution imaging presented in this manuscript is a valuable addition, since many cell biology studies performed on flat glass rely on the ability to image the distributions of different proteins at high precision and/or in 3D. This is particularly the case of mechanobiology approaches that aim to characterize cell adhesion, motility or signalling.

We have evidenced qualitative differences on cell behavior here. In this regard, the microstructured fractal substrates hold remarkable potential, as the fabrication process allows to vary between different lattice types and spacings as well as octahedra sizes and levels of detail, which can be used to evidence different regimes and the relevant parameters driving them. This work therefore paves the way for further characterization such as a screening of various proteins to determine the mechanisms underlying the adhesion of cells to the 3D substrates. Several studies were able to successfully elaborate a force generation model in adhesion structures through systematic protein colocalization using 3D SMLM, which evidenced their interaction at the nanoscale [62, 67, 63]. This, however, lies beyond the scope of this work, which reports a proof of concept of 3D SMLM imaging of cells cultivated on microstructured fractal substrates. Interestingly, the discrete nature of SMLM output data allows quantitative and statistical analyses such as determining local protein concentrations or assessing molecular clustering, spatial (co-)organization and redistribution. This will be relevant to explore in the near future to characterize the density and organization of focal adhesion complexes, which are essential parameters to understand the molecular mechanisms leading to cell differential behavior driven by physical cues. Although we limited this study to three days after seeding, longer growth times can also be interesting to investigate in order to determine the functional impact of the substrate. Finally, in order to determine sub-cellular functional changes induced by the culture on microstructured fractal substrates, live cell microscopy is often required. In this domain, SMLM is well suited since Single-Particle Tracking (SPT) allows the assessment of the distribution of diffusion coefficients of individual fluorescently-labelled proteins. This can reveal the interaction rates between different partners. SPT would be a valuable addition to cell biology studies, particularly for mechanobiology, where it could highlight the functional changes underlying structural behaviors observed on fixed cells, and our single-molecule localization proof of principle lead us to believe that SPT on the microstructured fractal substrates should be achievable.

## Acknowledgements

We acknowledge financial support by the French Agence Nationale de la Recherche, project ABC4M under the reference ANR-20-CE45-0023 (I.I. and C.C.), and project SP-Tunnel-OHG under the reference ANR-20-CE24-0021 (C.C.).

## Author contributions

C.C., M.C.C., A.S.A. and I.I. conceived the project. C.C. led its development with supervision from A.S.A. and I.I. and input from M.C.C. and E.B.. E.B. manufactured the microstructured fractal substrates, A.D.L. and K.P. prepared the biological samples from the cell culture to the fixation. C.C. performed the fluorescent labelling of both the microstructured fractal substrates and the cells. C.C. performed the white light and fluorescence imaging and E.B. performed the electron microscopy imaging. C.C. wrote the processing codes of the single-molecule localization microscopy data. C.C. wrote the first draft of the manuscript with input from M.C.C., E.B., S.L.G., A.S.A. and I.I. The manuscript was reviewed by all authors.

## Methods

### Microstructured fractal substrates fabrication

The fabrication involves depositing and pattering a hard oxide mask on the silicon wafer. Inverted pyramids are formed using anisotropic etching (25% KOH at 75°C). After RCA-2 cleaning, a selective etching step strips the SiO_2_ hard mask (1% HF), and the G0 mold is ready. G1–G3 is formed by a repetitive sequence of processing steps on top of substrate G0, which consists of dry oxidation at 1100°C with the formation of a conformal oxide formation over the silicon mold containing the topographic features. The SiO_2_ is conformally deposited, except at the sharp concave features of the mold. The concave features with the thinnest oxide thickness are selectively opened using timed isotropic etching (i.e., 1% HF etching). Through these openings, octahedral cavities bounded by Si (111) planes will form during a subsequent timed anisotropic etching (25% tetramethylammonium hydroxide (TMAH) at 70°C) step. This step determines the size of the octahedral features. G1 is tuned so that the largest width equals the size of the base of the mold G0. Repeating the sequence described above and reducing the etching time by half compared to the previous step will decrease the feature size by a factor 2, while the number of octahedral features increases by a factor 5. This way, G2 and G3 are formed [26, 28]. In the case of G4, the final iteration on top of G3 is, instead of relying on oxidation retardation, performed by the following processing steps: silicon nitride corner lithography, local oxidation of silicon, and selective stripping of the remaining silicon nitride. The next step is the same: timed anisotropic etching (25% TMAH at 70°C) step [26, 28]. After creating all G0–G4 mold wafers, a structural SiO_2_ thin film is grown, and these wafers are anodically bonded to borosilicate glass wafers. By pre-dicing in 1 cm^2^ pieces and dissolving the silicon mold wafer in the anisotropic etchant with high selectivity against SiO_2_ (25% TMAH at 70°C), the 1 cm^2^ topographic substrates are ready to use [26, 28].

### Functionalized microstructured fractal substrates preparation

The substrates were labelled with N-Hydroxysuccinimide (NHS) ester-functionalized Abberior Cage 635 dyes using the following protocol. First, 25 µl of 0.1 % Bovine Serum Albumin (BSA) (Sigma, A9576) in Phosphate Buffered Saline (PBS) were added. After 15 minutes, the substrate was washed with PBS. Then, a solution containing 40 µl of PBS (pH 7.4), 5 µl of sodium bicarbonate 7.5 % (Sigma, S8761) (pH 8.0) and 1 µl of NHS ester Abberior Cage 635 was prepared and added to the substrate. After 10 minutes, the substrate was washed with PBS.

### Cell culture, fixation and fluorescent labelling

Human mesenchymal stem cells (hMSCs) (ATCC, lgc-standards, Teddington, UK) were cultured in high-glucose DMEM (Gibco) supplemented with 10% FBS and 1 % penicillin/streptomycin. Control differentiation experiments were conducted using osteogenic differentiation medium (Sigma, C-28013) and adipogenic differentiation medium (Sigma, C-28016) following the manufacturers’ instructions.

Flat glass and microstructured fractal substrates were meticulously cleaned with ethanol, dried, and washed with PBS before cell seeding. For differentiation experiments hMSCs were seeded at a density of 20,000 cells per cm^2^ on the prepared surfaces in a 24-well plate. Subsequently, 1 ml of standard medium was added to the cells, for control differentiation experiments on glass surfaces 1 ml of the respective differentiation media were used. Media were refreshed every three days. After 14 days, cells were washed with PBS and fixated using 4 % paraformaldehyde in PBS for 10 minutes at room temperature. The fixative was then removed, and the cells were extensively washed with PBS for further analysis. Osteogenic differentiation was assessed by alkaline phosphatase staining (Sigma, B5655) as per the manufacturers’ guidelines. Adipogenic differentiation was evaluated using an oil red O assay (Sigma, O0625) following the manufacturers’ instructions.

For SMLM, hMSCs were seeded on meticulously cleaned flat glass and microstructured fractal substrates at a density of 10,000 cells per cm^2^ in a 24-well plate. Subsequently, 1 mL of standard medium was added to the cells. After 3 days, cells were washed with PBS and fixated using 4 % paraformaldehyde in PBS for 10 minutes at room temperature. The fixative was then removed, and the cells were extensively washed with PBS.

The actin labelling of the fixed cells was performed by adding Alexa Fluor 647 phalloidin (Thermo Fisher, A22287) at a concentration of 10 nM in the imaging buffer (PBS or dSTORM buffer for wide field fluorescence and SMLM respectively, see the corresponding sections of the **Methods**). The imaging was performed immediately after 20 min of staining without washing.

### SEM imaging of the microstructured fractal substrates

The SEM images were obtained using a JEOL JSM 7610FPlus FEG SEM system equipped with Inlens and High-Efficiency Secondary Electrons. Acceleration voltages between 1 kV and 5 kV were applied for the SEM image collection.

### Optical setup

A general schematic of the optical setup used is presented in **Supplementary Fig. 1a**. Depending on the imaging modality, different microscopy setups were used. For the SMLM acquisitions and the wide field fluorescence images in **Fig. 1e**, we used a custom-built inverted microscope with a RM21 body and a MANNZ micro- and nano-positioner (Mad City Labs). The illumination and fluorescence collection was done with a Nikon 100x 1.49NA APO TIRF SR oil immersion objective. The excitation was performed thanks to a 638 nm laser (LBX-638-180, 180 mW, Oxxius) with a 405 nm laser for pumping (LBX-405-50, 50 mW, Oxxius). A multiband filter set (LF405/488/561/635-A-000, Semrock) was used. The excitation consisted of a standard vertical Gaussian beam. The fluorescence was recorded on an EMCCD camera (iXon Ultra 897, Andor). We used an afocal doublet to adjust the pixel size to 107 nm in the object plane. A cylindrical lens of focal length 700 mm (Thorlabs, LJ1836L1-A) was added to create the astigmatism.

The white light transmission images and wide-field diffraction-limited fluorescence images in **Fig. 4** and **Supplementary Video 5** were performed on an inverted Ti2 Eclipse (Nikon) microscope equipped with a Nano-Z nanopositioner (Mad City Labs) and an Olympus Plan N 10x 0.25NA air objective (for the white light transmission images, as well as for **Fig. 4**) or a Nikon Plan Apo *λ* 60x 1.4NA oil immersion objective (for **Supplementary Video 5**). A white lamp was used for the transmission images. For the wide field fluorescence imaging, the excitation consisted of a vertical Gaussian beam produced by a 647 nm laser (LBX-647-140, 140 mW, Oxxius). A multiband filter set (LF405/488/561/635-A-000, Semrock) was used. The image was formed on an sCMOS camera (Prime 95B, Photometrics). We used an afocal doublet to adjust the pixel size to 660 nm and 110 nm in the object plane for the 10x and the 60x objectives respectively.

### Sample mounting

As the thickness of the substrate (∼500µm) is larger than the working distance (∼120 µm) of the high numerical aperture objective used for the SMLM acquisitions, imaging through the substrate was not possible. Therefore, the microstructured fractal substrates were flipped down and deposited on a 150 µm thick coverslip (Marienfeld, 0111640, 24 mm diameter) containing the imaging medium. More precisely, we used an Attofluor chamber (Thermo Fisher, A7816) mounted with the 150-µm coverslip. Then we filled the chamber with a large volume (500 µl) of imaging buffer to avoid evaportaion effects, and we immersed the microstructured fractal substrate facing down. Finally, using tweezers, we gently squeezed the substrate onto the coverslip to reduce the spacing and avoid motion of the substrate during imaging. Although the chamber was not sealed, the sample remained stable during the course of the acquisition (20–40 minutes). We found that the tips of the microstructures were not directly in contact with the bottom coverslip, but rather located approximately one to several µm above, which is likely due to the curvature of the bottom coverslip (in other words, the fractals are in contact at the edges of the substrate only). The general sample mounting geometry is illustrated in **Fig. 1d**. Note that 170 µm thick coverslips can also be used as an alternative (for similar results), as the working distance of the objective is sufficient to focus on the microstructured fractal substrates given the low spacing between the coverslip and the substrate.

For the substrates with perforated tips (**Fig. 3c–d**, **Supplementary Fig. 3**), we first filled the substrates during 5 minutes with pure ethanol, which exhibits lower surface tension than aqueous buffers. Therefore, the ethanol quickly flowed through the nanometric holes to fill the octahedra. Then, the substrate was washed with PBS several times and we added the imaging buffer to replace the ethanol.

### Diffraction-limited fluorescence imaging

The samples were illuminated with a 647 nm continuous excitation at an irradiance of 0.1 kW/cm^2^. The sCMOS camera exposure time was set to 100 ms. For the *z* stacks, the sample was moved using the piezoelectric stage. The microstructured fractal substrates functionalized with Abberior Cage 635 were imaged in PBS, while the actin-stained cells were imaged in PBS containing the Alexa Fluor 647 (see **Cell culture, fixation and fluorescent labelling**).

### Single-molecule imaging of the functionalized microstructured fractal substrates

The samples were imaged in PBS. They were illuminated with a 638 nm continuous excitation at an irradiance of 5 kW/cm^2^. The EMCCD gain and exposure time were set to 100 and 30 ms respectively. A low-power continuous 405 nm laser beam was used to photoactivate the dyes.

For the extended 3D imaging of the functionalized microstructured fractal substrates, the piezoelectric stage was used to move the focal plane by 100 nm every 30 seconds. Taking into account the depth of field (∼1 µm), this implies that each plane *z* position remained in the imaging range during 10000 frames. Localizing the axial position of each molecule relative to the focal plane and translating it into an absolute axial position thanks to the known position of the piezoelectric stage at each time enables the reconstruction of the 3D shape of the whole structures.

### Single-molecule imaging of the biological samples

The samples were imaged in a dSTORM buffer containing 100 mg/ml glucose, 3.86 mg/ml MEA, 0.5 mg/ml glucose oxidase and 1.18 µl/ml catalase in PBS, as well as the Alexa Fluor 647 phalloidin (see **Cell culture, fixation and fluorescent labelling**). They were illuminated with a 638 nm continuous excitation at an irradiance of 2 kW/cm^2^. The EMCCD gain and exposure time were set to 50 and 50 ms respectively. A low-power continuous 405 nm laser beam was used to pump the molecules from the dark state back to the fundamental state and therefore increase the PSF density.

### Axial calibration

40-nm fluroescent beads (F10720, Thermo Fisher) were deposited on a substrate with a dilution factor of 1:10^7^ in phosphate buffered saline (PBS). After waiting a few minutes for the beads to deposit, a sufficient density of beads was achieved. The solution was removed and the substrate was flipped down on a 150 µm thick coverslip with PBS as imaging buffer. This corresponds to a sample mounting geometry identical to that used for the samples of interest (see the **Sample mounting** section in the **Methods**).

The piezoelectric stage was used to acquire a *z* stack by moving the sample by 100-nm steps. Each bead in the *z* stack was localized to determine its *x* and *y* widths, written *w_x_* and *w_y_*respectively. By taking into account the value of the focal shift (0.75, see **Fig. 3e**), the curve that links *w_x_* − *w_y_*with the *z* position relative to the focal plane was generated for each bead. The astigmatism calibration curve was taken as the average of the individual curves for each bead.

### Localization and data treatment software

All the processing and rendering was performed using a home-written Python code. The temporal stack of raw frames was first filtered in a sliding window manner: for each frame *i*, the pixel-per-pixel median of the frames [*i* −100, *i* +100] was subtracted. This allows to remove the static or slowly varying background while keeping the PSFs unchanged. Then, on each frame, the PSFs were detected using a wavelet filtering algorithm [68], and localized by fitting an anisotropic Gaussian (i.e. allowing different widths in *x* and *y*). Finally, the lateral drift was corrected either using fiducial markers or from the localized molecules using a Direct Cross-Correlation (DCC) algorithm. The localization code is available at the following address: https://github.com/Clement-Cabriel/Frame-based-SMLM

## Data availability

Dataset are available from the corresponding author upon reasonable request.

## Supplementary material

**Supplementary Figure 1:**
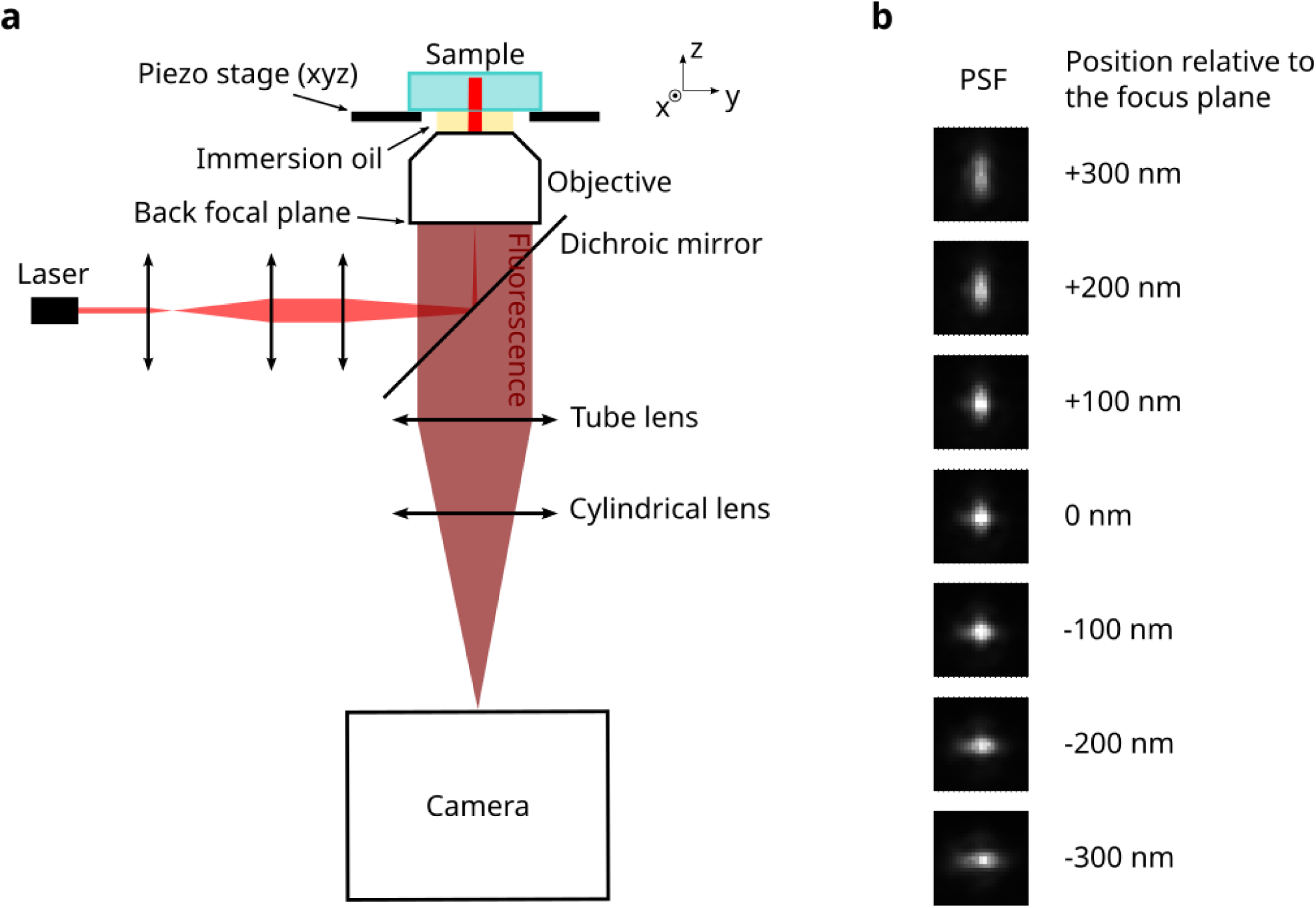
Optical setup and implementation of 3D SMLM. **a** Optical setup. The excitation consists of a simple collimated Gaussian laser beam. The fluorescence is collected by the same objective and separated from the excitation by a dichroic mirror. A tube lens forms the image on the camera and a cylindrical lens induces the astigmatism used for the 3D detection. Note that for the 2D SMLM, the setup is almost identical, the only difference being that there is no cylindrical lens for the 2D SMLM. **b** Illustration of the deformation of the PSF induced by the astigmatism as a function of the axial position of the molecule relative to the focus plane. The total width of the PSF images displayed is 2.7 µm.

**Supplementary Figure 2:**
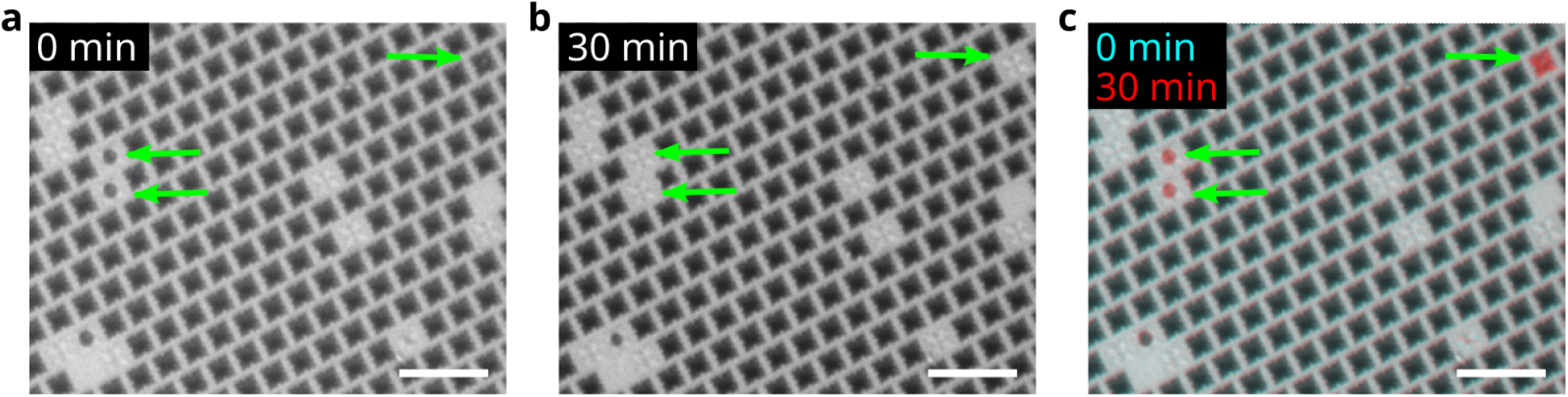
White light transmission imaging of the spontaneous filling of fractal structures by the imaging buffer. Two images of the same field of view are acquired separated by 30 minutes (**a–b**). The arrows indicate the few fractal structures that spontaneously fill with imaging buffer. The overlay of the two images is also displayed for clarity (**c**). Scale bars: 40 µm.

**Supplementary Figure 3:**
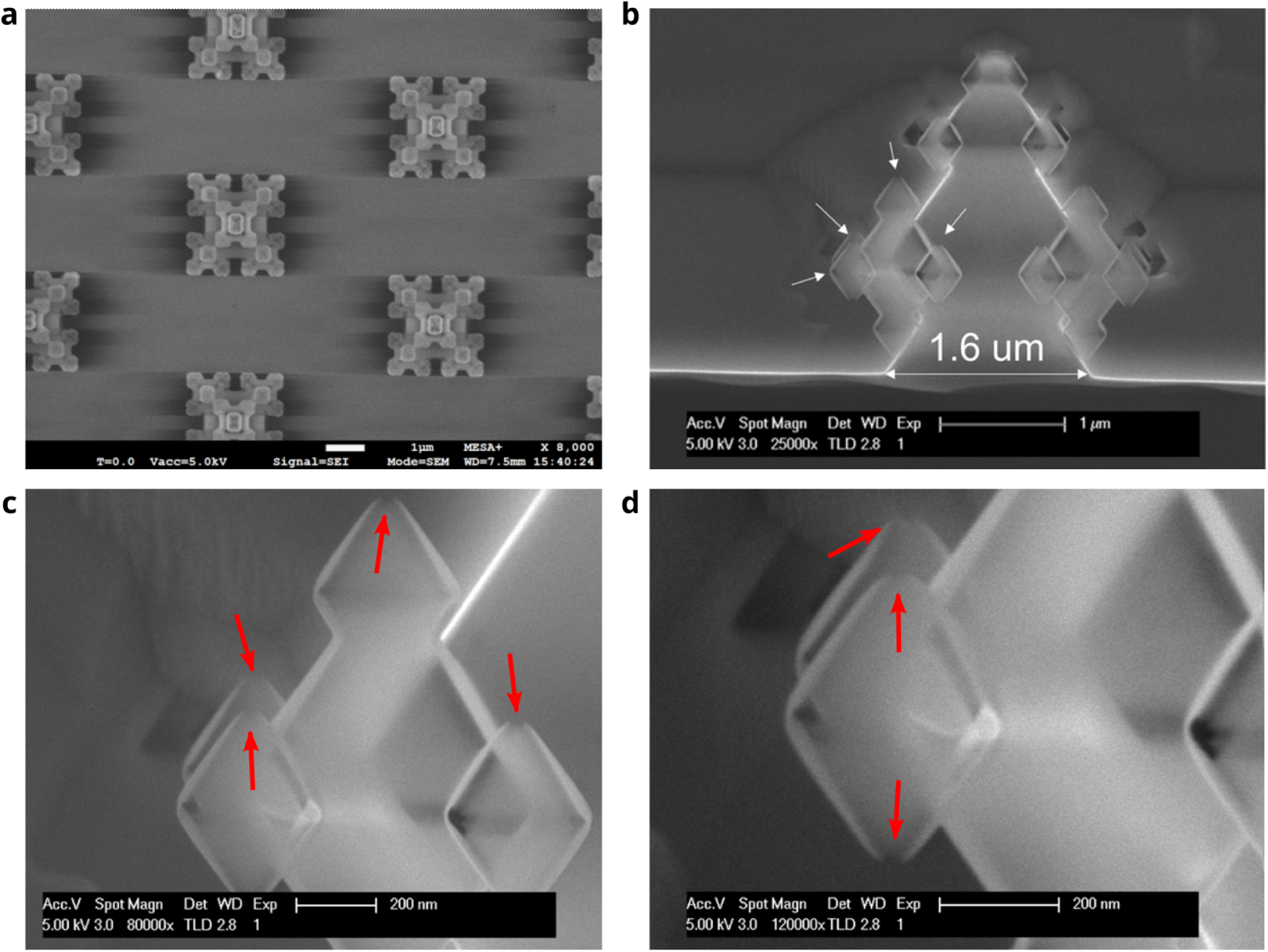
SEM imaging of the perforated G3 microstructured fractal substrates used in **Fig. 3c–d**. **a** Top view showing the general geometry. **b–d** Side views at different magnification levels showing the holes at the tips (indicated with the arrows). Scale bars: 1 µm (**a–b**), 200 nm (**c–d**).

**Supplementary Figure 4:**
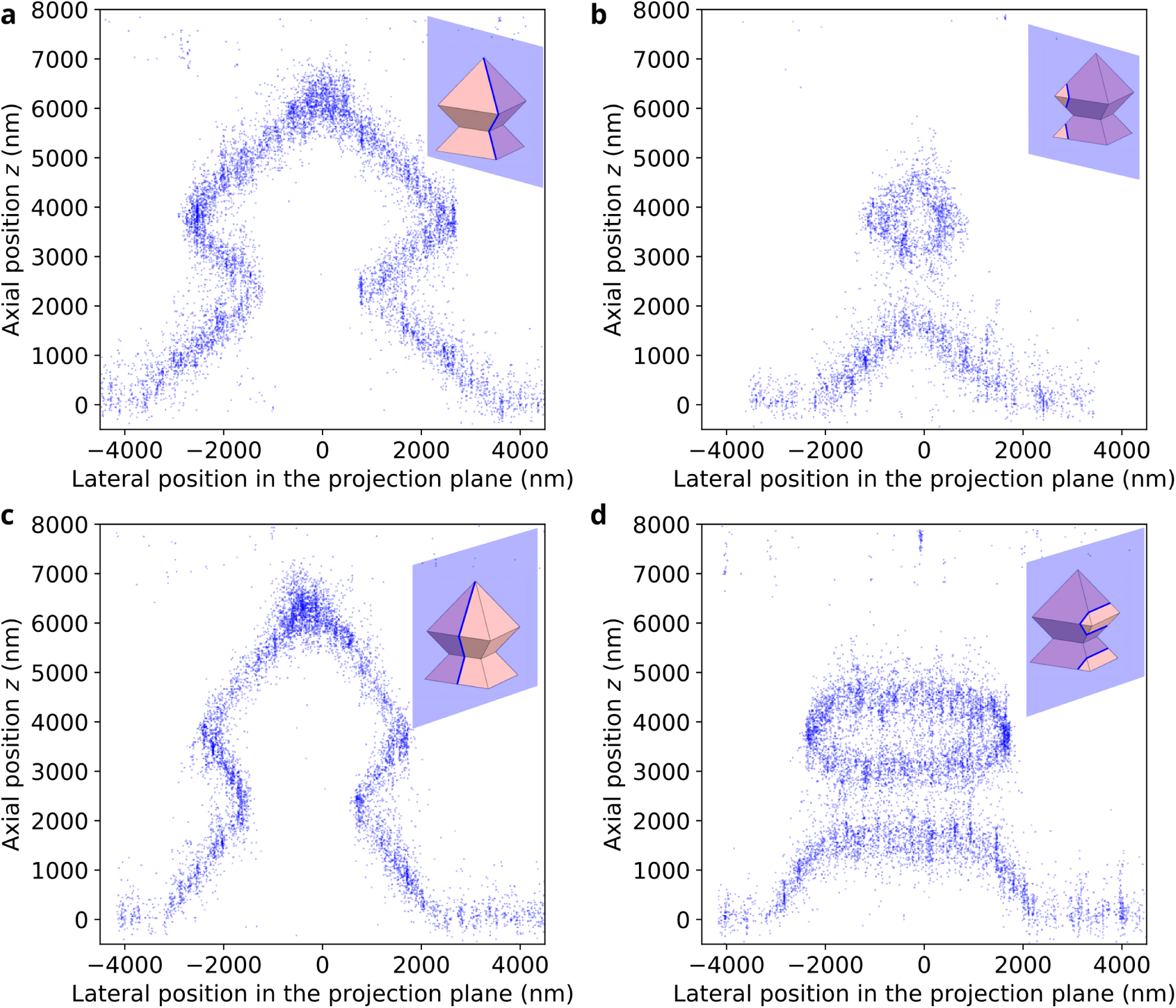
3D slices of the fluorescently-labelled G1 microstructure imaged in 3D SMLM (similar to Fig. 3e). Thin slices (500-nm thick) are generated by post-processing the 3D SMLM results, and the molecules are projected in two different planes—parallel to the diagonal (**a,b**) or parallel to the sides (**c,d**). Various positions are also investigated—centered on the fractal structure (**a,c**) or off-center (**b,d**). The corresponding planes are displayed in the insets. This type of representations is straightforward to generate in 3D SMLM, unlike in electron microscopy.

### Supplementary Note 1: Comparative SEM and SMLM distance measurements

Lateral distances of the different fractal features are measured from both the 2D SMLM images (acquired in the middle plane) and the SEM images (top view). As a convention, distance measurements always refer to the side of the octahedra rather than their diagonal unless otherwise stated. The measurements are performed in both directions, i.e. *x* and *y*. This is illustrated in **Supplementary Fig. 5**. For each sample, we measure all the features up to the number of iterations of the sample—for example, G3 samples allow the measurements of the G1, G2 and G3 feature sizes. Additionally, we measure the spacing between adjacent G4 octahedra, as defined in Fig. 2c. The results are presented in **Supplementary Table 1**, which displays the mean values and standard deviations measured over a statistical set of measurements. It should be noted that the measured structures are not exactly the same (as it is the case in Correlative Light Electron Microscopy), which can explain the differences in the standard deviation values displayed.

**Supplementary Figure 5:**
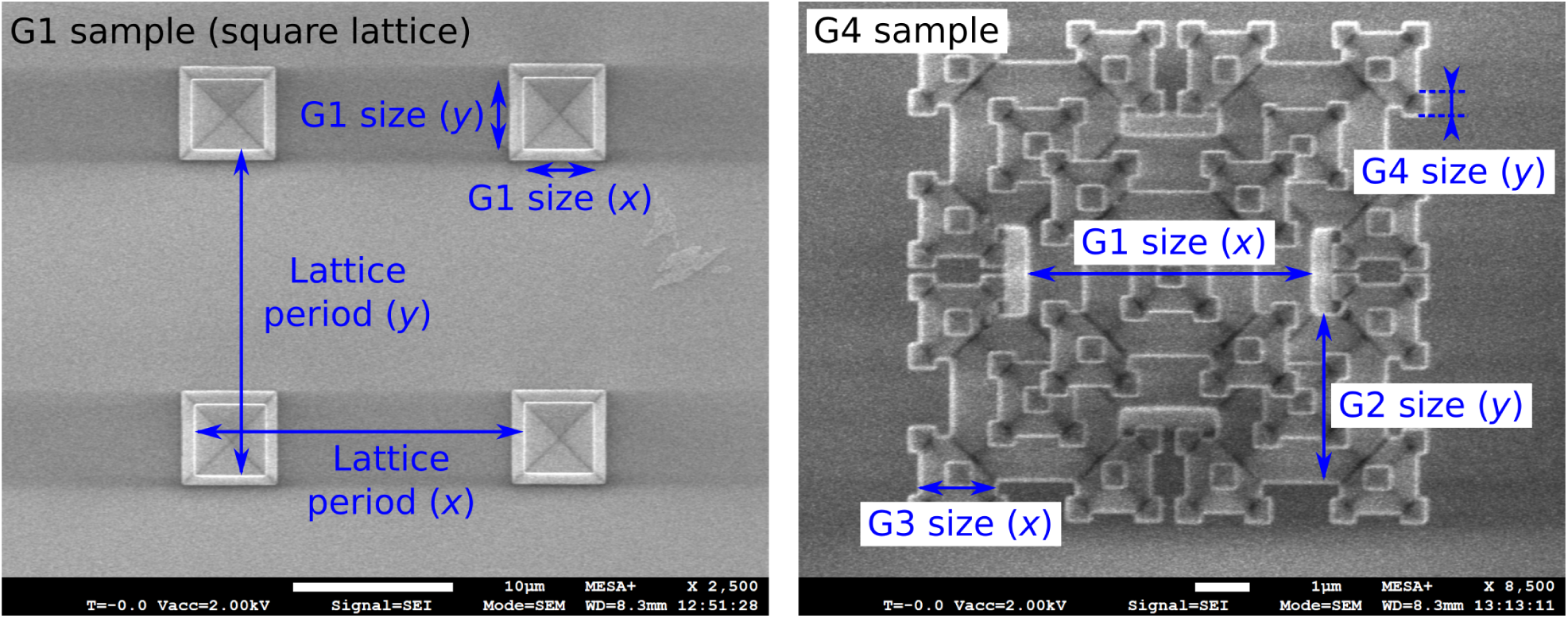
Principle of the measurements of the different fractal features in SEM. Left: G1 sample (square lattice) showing the measurements of the lattice period and the G1 features in *x* and *y*. Right: G4 sample showing the measurements of the G1, G2, G3 and G4 features in *x* and *y*. The same principle is applied for the SMLM measurements. Scale bars: 10 µm (left), 1 µm (right).

**Supplementary Table 1:** Summary of the different distance measurements performed on G1, G3 and G4 fractal structures (square lattice). The distance values displayed are the mean ± 2 × standard deviation of the distribution, and the size of the statistical set is indicated in parentheses.

| Sample | Corresponding figure | Measured features | Sizes measured in SEM (nm) | Sizes measured in SMLM (nm) |
| --- | --- | --- | --- | --- |
| G1, square | **Fig. 2a** | Lattice period, $x$<br>Lattice period, $y$<br>G1, $x$<br>G1, $y$ | $20660 \pm 260$ (2)<br>$20260 \pm 320$ (2)<br>$4390 \pm 120$ (6)<br>$4410 \pm 140$ (6) | $20360 \pm 250$ (2)<br>$20300 \pm 310$ (2)<br>$4340 \pm 240$ (6)<br>$4320 \pm 120$ (6) |
| G3, square | **Fig. 2b top** | G1, $x$<br>G1, $y$<br>G2, $x$<br>G2, $y$<br>G3, $x$<br>G3, $y$ | $5358 \pm 120$ (6)<br>$5205 \pm 86$ (6)<br>$3040 \pm 30$ (18)<br>$2960 \pm 42$ (18)<br>$1330 \pm 28$ (21)<br>$1299 \pm 29$ (21) | $5360 \pm 80$ (4)<br>$5250 \pm 90$ (4)<br>$3022 \pm 68$ (13)<br>$2985 \pm 69$ (13)<br>$1327 \pm 72$ (23)<br>$1295 \pm 74$ (23) |
| G4, square | **Fig. 2b bottom** | G1, $x$<br>G1, $y$<br>G2, $x$<br>G2, $y$<br>G3, $x$<br>G3, $y$<br>G4, $x$<br>G4, $y$ | $5206 \pm 64$ (5)<br>$5100 \pm 60$ (5)<br>$3124 \pm 47$ (12)<br>$3062 \pm 33$ (12)<br>$1381 \pm 25$ (22)<br>$1360 \pm 15$ (22)<br>$457 \pm 22$ (18)<br>$448 \pm 20$ (18) | $5213 \pm 60$ (3)<br>$5100 \pm 50$ (3)<br>$3133 \pm 76$ (10)<br>$3078 \pm 73$ (10)<br>$1394 \pm 51$ (25)<br>$1361 \pm 48$ (25)<br>$453 \pm 26$ (33)<br>$449 \pm 29$ (33) |
| G4, square | **Fig. 2c** | G4 spacing, $x$<br>G4 spacing, $y$ | $205 \pm 43$ (4)<br>$110 \pm 52$ (4) | $180 \pm 17$ (4)<br>$103 \pm 21$ (4) |

### Supplementary Video 1: Extracts of the blinking movies acquired at the three planes shown in Fig. 3a

The video presents extracts from the blinking movie acquired slightly above the middle plane (left), slightly below the middle plane (center) and slightly above the flat bottom substrate (right). The imaging planes correspond to those imaged in Fig. 3a and illustrated in the insets. The total height of the field (in *y*) is 34 µm and the movie is played at the real acquisition speed (33 frames per second).

Note that for the fractal structure on the top left (i.e. that indicated with a cyan dotted square in Fig. 3a), the PSFs are clearly visible in the three imaging planes, while for the one at the bottom left (magenta dotted square), they are clearly visible only above the middle plane, and are strongly distorted to invisible in the other two planes.

### Supplementary Video 2: Extracts of the blinking movie acquired with the perforated fractal structures

The video presents extracts from the blinking movies acquired slightly above the flat bottom substrate with the perforated G3 fractal structures (right) labelled with fluorescent molecules (see Fig. 3d). The left panel shows the blinking movie obtained in the same conditions with the non-perforated fractal structures (i.e. the same data as in **Supplementary Video 1 right**) as a comparison—note that the lattice periods and octahedra sizes are not the same in both cases though. The total height of the fields (in *y*) is 14 µm and the movie is played at the real acquisition speed (33 frames per second).

Note that the PSFs are clearly visible for all the perforated fractal structures (right)—on the contrary, there are no visible PSFs close to the substrate under the fractal structure filled with air (left).

### Supplementary Video 3: 3D visualization of the fluorescently-labelled G3 fractal structure presented in Fig. 3h

The video is an animation of the static representation presented in Fig. 3h. The molecules detected on the labelled G3 fractal structure are displayed as a projection on a given plane, which is rotated by increments of 5°, first horizontally (first half of the video) and then vertically (second half). The rotation angles along the two axes are indicated on each frame. The total width and height of the video correspond to 13 µm each, and the color encodes the position along the axis orthogonal to the projection plane.

It should be noted that this type of representation is straightforward to obtain since the output of 3D SMLM is the 3D position of each molecule detected, therefore generating projections along various planes only requires the proper display code. On the contrary, electron microscopy experiments return information which is already on a plane selected during the acquisition, making it difficult to obtain a real 3D representation.

### Supplementary Video 4: Wide field microscopy ***z*** stacks of cells cultivated on flat glass substrates

The video is a set of two *z* stacks of hMSCs cultivated on flat glass substrates, labelled with phalloidin- Alexa Fluor 647 (targetting the F-actin) and imaged in wide-field diffraction-limited microscopy, similar to Fig. 4. Fluorescent beads were also added to the sample. Each step corresponds to an axial motion of the piezoelectric stage of 500 nm, i.e. a displacement of the focal plane of 375 nm taking into account the focal shift (0.75). The *z* position is indicated in each frame. The scan is performed from the top (towards the imaging medium) to the bottom (substrate). The total height (in *y*) of the fields displayed is 114 µm. The planes are displayed at a rate of two per second (i.e. 750 nm per second), for a total axial range (in *z*) of 19.5 µm.

Note that the dense regions shown here, cells grow as multiple layers resting on top of each other, without any preferential direction.

### Supplementary Video 5: Wide field microscopy ***z*** stacks of cells cultivated on microstructured fractal substrates

The video is a set of three *z* stacks of hMSCs cultivated on square lattice G3 microstructured fractal substrates, labelled with phalloidin-Alexa Fluor 647 (targetting the F-actin) and imaged in wide-field diffraction-limited microscopy, similar to Fig. 4. Each step corresponds to an axial motion of the piezoelectric stage of 500 nm, i.e. a displacement of the focal plane of 375 nm taking into account the focal shift (0.75). The *z* position is indicated in each frame. The scan is performed from the top (tips of the octahedra) to the bottom (flat part of the substrate). The total height (in *y*) of the fields displayed is 114 µm. The planes are displayed at a rate of two per second (i.e. 750 nm per second), for a total axial range (in *z*) of 11 µm. Three different fields are displayed (left, center and right).

The region on the left shows a cell mostly resting on the base substrate and growing in the grooves between the fractal structures, while the at the center displays a cell growing mostly in the middle axial plane of the microstructures, and using them as anchor points to remain suspended above the base substrate. Finally, the region on the right corresponds to a higher density, where the cells tend to grow as one or several flat layers resting on the upper tips of the octahedra, thus exhibiting a rather isotropic behavior, i.e. the actin fibers do not seem to significantly align with the axes of the lattice.

## References

[1] A. M. Ross, Z. Jiang, M. Bastmeyer, and J. Lahann, “Physical aspects of cell culture substrates: Topography, roughness, and elasticity,” Small, vol. 8, pp. 336–355, Dec. 2011.

[2] K. Anselme and M. Bigerelle, “Role of materials surface topography on mammalian cell response,” International Materials Reviews, vol. 56, pp. 243–266, July 2011.

[3] K. Metavarayuth, P. Sitasuwan, X. Zhao, Y. Lin, and Q. Wang, “Influence of surface topographical cues on the differentiation of mesenchymal stem cells in vitro,” ACS Biomaterials Science & Engineering, vol. 2, pp. 142–151, jan 2016.

[4] M. Ferrari, F. Cirisano, and M. C. Morán, “Mammalian cell behavior on hydrophobic substrates: Influence of surface properties,” Colloids and Interfaces, vol. 3, p. 48, May 2019.

[5] N. Monteiro, J. Fangueiro, R. Reis, and N. Neves, “Replication of natural surface topographies to generate advanced cell culture substrates,” Bioactive Materials, vol. 28, pp. 337–347, Oct. 2023.

[6] E. Fennema, N. Rivron, J. Rouwkema, C. van Blitterswijk, and J. de Boer, “Spheroid culture as a tool for creating 3d complex tissues,” Trends in Biotechnology, vol. 31, pp. 108–115, feb 2013.

[7] N. Picollet-D’hahan, M. E. Dolega, L. Liguori, C. Marquette, S. Le Gac, X. Gidrol, and D. K. Martin, “A 3d toolbox to enhance physiological relevance of human tissue models,” Trends in Biotechnology, vol. 34, pp. 757–769, sep 2016.

[8] N.-E. Ryu, S.-H. Lee, and H. Park, “Spheroid culture system methods and applications for mesenchymal stem cells,” Cells, vol. 8, p. 1620, dec 2019.

[9] Z. Zhao, X. Chen, A. M. Dowbaj, A. Sljukic, K. Bratlie, L. Lin, E. L. S. Fong, G. M. Balachander, Z. Chen, A. Soragni, M. Huch, Y. A. Zeng, Q. Wang, and H. Yu, “Organoids,” Nature Reviews Methods Primers, vol. 2, dec 2022.

[10] Y. Torisawa, B. Mosadegh, G. D. Luker, M. Morell, K. S. O’Shea, and S. Takayama, “Microfluidic hydrodynamic cellular patterning for systematic formation of co-culture spheroids,” Integrative Biology, vol. 1, no. 11-12, p. 649, 2009.

[11] A. Sridhar, H. L. de Boer, A. van den Berg, and S. Le Gac, “Microstamped petri dishes for scanning electrochemical microscopy analysis of arrays of microtissues,” PLoS ONE, vol. 9, p. e93618, apr 2014.

[12] H. Geckil, F. Xu, X. Zhang, S. Moon, and U. Demirci, “Engineering hydrogels as extracellular matrix mimics,” Nanomedicine, vol. 5, pp. 469–484, apr 2010.

[13] M. W. Tibbitt and K. S. Anseth, “Hydrogels as extracellular matrix mimics for 3d cell culture,” Biotechnology and Bioengineering, vol. 103, pp. 655–663, may 2009.

[14] Y. Zhang, Y. Xu, and J. Gao, “The engineering and application of extracellular matrix hydrogels: a review,” Biomaterials Science, vol. 11, no. 11, pp. 3784–3799, 2023.

[15] M. Jin, G. Koçer, and J. I. Paez, “Luciferin-bioinspired click ligation enables hydrogel platforms with fine-tunable properties for 3d cell culture,” ACS Applied Materials & Interfaces, vol. 14, pp. 5017–5032, jan 2022.

[16] N. C. Rivron, E. J. Vrij, J. Rouwkema, S. Le Gac, A. van den Berg, R. K. Truckenmüller, and C. A. van Blitterswijk, “Tissue deformation spatially modulates VEGF signaling and angiogenesis,” Proceedings of the National Academy of Sciences, vol. 109, pp. 6886–6891, apr 2012.

[17] A. Clancy, D. Chen, J. Bruns, J. Nadella, S. Stealey, Y. Zhang, A. Timperman, and S. P. Zustiak, “Hydrogel-based microfluidic device with multiplexed 3d in vitro cell culture,” Scientific Reports, vol. 12, oct 2022.

[18] F. Dituri, M. Centonze, E. Berenschot, N. R. Tas, A. Susarrey-Arce, and S. Krol, “Complex tumor spheroid formation and one-step cancer-associated fibroblasts purification from hepatocellular carcinoma tissue promoted by inorganic surface topography,” Nanomaterials, vol. 11, p. 3233, nov 2021.

[19] P. J. Amos, E. C. Bozkulak, and Y. Qyang, “Methods of cell purification: A critical juncture for laboratory research and translational science,” Cells Tissues Organs, vol. 195, pp. 26–40, oct 2011.

[20] M. Centonze, E. Berenschot, S. Serrati, A. Susarrey-Arce, and S. Krol, “The fast track for intestinal tumor cell differentiation and in vitro intestinal models by inorganic topographic surfaces,” Pharmaceutics, vol. 14, p. 218, jan 2022.

[21] S. C. Carrara, A. Dávila-Lezama, C. Cabriel, E. Berenschot, S. Krol, H. Gardeniers, I. Izeddin, H. Kolmar, and A. Susarrey-Arce, “3D topographies promote macrophage M2d-subset differentiation,” Materials Today Bio, p. 100897, Dec. 2023.

[22] A. Curtis and C. Wilkinson, “Nantotechniques and approaches in biotechnology,” Trends in Biotechnology, vol. 19, pp. 97–101, mar 2001.

[23] L. E. McNamara, T. Sjöström, K. E. Burgess, J. J. Kim, E. Liu, S. Gordonov, P. V. Moghe, R. D. Meek, R. O. Oreffo, B. Su, and M. J. Dalby, “Skeletal stem cell physiology on functionally distinct titania nanotopographies,” Biomaterials, vol. 32, pp. 7403–7410, oct 2011.

[24] M. J. Dalby, N. Gadegaard, R. Tare, A. Andar, M. O. Riehle, P. Herzyk, C. D. W. Wilkinson, and R. O. C. Oreffo, “The control of human mesenchymal cell differentiation using nanoscale symmetry and disorder,” Nature Materials, vol. 6, pp. 997–1003, sep 2007.

[25] A. R. Roy, W. Zhang, Z. Jahed, C.-T. Tsai, B. Cui, and W. E. Moerner, “Exploring cell surface–nanopillar interactions with 3d super-resolution microscopy,” ACS Nano, vol. 16, pp. 192– 210, sep 2021.

[26] E. Berenschot, H. V. Jansen, and N. R. Tas, “Fabrication of 3d fractal structures using nanoscale anisotropic etching of single crystalline silicon,” Journal of Micromechanics and Microengineering, vol. 23, p. 055024, apr 2013.

[27] D. Jonker, C. Eyovge, E. Berenschot, V. Di Palma, D. Wasserberg, S. Michel-Souzy, P. Jonkheijm, S. Krol, H. Gardeniers, M. Creatore, N. Tas, and A. Susarrey-Arce, “Electrochemical sensing with spatially patterned pt octahedra electrodes,” Advanced Materials Technologies, vol. 9, Jan. 2024.

[28] E. Berenschot, R. Tiggelaar, J. Geerlings, H. Gardeniers, N. Tas, M. Malankowska, M. Pina, and R. Mallada, “3d-fractal engineering based on oxide-only corner lithography,” in 2016 Symposium on Design, Test, Integration and Packaging of MEMS/MOEMS (DTIP), IEEE, may 2016.

[29] I. Izeddin, M. El Beheiry, J. Andilla, D. Ciepielewski, X. Darzacq, and M. Dahan, “PSF shaping using adaptive optics for three-dimensional single-molecule super-resolution imaging and tracking.,” Optics express, vol. 20, pp. 4957–67, feb 2012.

[30] D. Burke, B. Patton, F. Huang, J. Bewersdorf, and M. J. Booth, “Adaptive optics correction of specimen-induced aberrations in single-molecule switching microscopy,” Optica, vol. 2, p. 177, feb 2015.

[31] Y. Deng and J. W. Shaevitz, “Effect of aberration on height calibration in three-dimensional localization-based microscopy and particle tracking,” Applied Optics, vol. 48, p. 1886, mar 2009.

[32] R. McGorty, J. Schnitzbauer, W. Zhang, and B. Huang, “Correction of depth-dependent aberrations in 3d single-molecule localization and super-resolution microscopy,” Optics Letters, vol. 39, p. 275, jan 2014.

[33] R. Tafteh, D. R. L. Scriven, E. D. W. Moore, and K. C. Chou, “Single molecule localization deep within thick cells; a novel super-resolution microscope,” Journal of Biophotonics, vol. 9, pp. 155– 160, aug 2015.

[34] C. Cabriel, N. Bourg, G. Dupuis, and S. Lévêque-Fort, “Aberration-accounting calibration for 3D single-molecule localization microscopy,” Optics Letters, vol. 43, p. 174, jan 2018.

[35] Y. Li, Y.-L. Wu, P. Hoess, M. Mund, and J. Ries, “Depth-dependent PSF calibration and aberration correction for 3d single-molecule localization,” Biomedical Optics Express, vol. 10, p. 2708, may 2019.

[36] P. N. Petrov and W. E. Moerner, “Addressing systematic errors in axial distance measurements in single-emitter localization microscopy,” Optics Express, vol. 28, p. 18616, jun 2020.

[37] E. Betzig, G. H. Patterson, R. Sougrat, O. W. Lindwasser, S. Olenych, J. S. Bonifacino, M. W. Davidson, J. Lippincott-Schwartz, and H. F. Hess, “Imaging intracellular fluorescent proteins at nanometer resolution,” Science, vol. 313, pp. 1642–1645, Sept. 2006.

[38] S. T. Hess, T. P. K. Girirajan, and M. D. Mason, “Ultra-high resolution imaging by fluorescence photoactivation localization microscopy,” Biophysical journal, vol. 91, pp. 4258–72, dec 2006.

[39] M. J. Rust, M. Bates, and X. Zhuang, “Sub-diffraction-limit imaging by stochastic optical reconstruction microscopy (STORM),” Nature Methods, vol. 3, pp. 793–796, aug 2006.

[40] B. Huang, W. Wang, M. Bates, and X. Zhuang, “Three-Dimensional Super-Resolution Imaging by Stochastic Optical Reconstruction Microscopy,” Science, vol. 319, no. 5864, pp. 810–813, 2008.

[41] M. F. Juette, T. J. Gould, M. D. Lessard, M. J. Mlodzianoski, B. S. Nagpure, B. T. Bennett, S. T. Hess, and J. Bewersdorf, “Three-dimensional sub-100 nm resolution fluorescence microscopy of thick samples.,” Nature methods, vol. 5, no. 6, pp. 527–529, 2008.

[42] S. R. P. Pavani, J. G. DeLuca, and R. Piestun, “Polarization sensitive, three-dimensional, single-molecule imaging of cells with a double-helix system.,” Optics express, vol. 17, no. 22, pp. 19644– 19655, 2009.

[43] G. Shtengel, J. A. Galbraith, C. G. Galbraith, J. Lippincott-Schwartz, J. M. Gillette, S. Manley, R. Sougrat, C. M. Waterman, P. Kanchanawong, M. W. Davidson, R. D. Fetter, and H. F. Hess, “Interferometric fluorescent super-resolution microscopy resolves 3d cellular ultrastructure,” Proceedings of the National Academy of Sciences, vol. 106, pp. 3125–3130, feb 2009.

[44] P. Bon, J. Linarès-Loyez, M. Feyeux, K. Alessandri, B. Lounis, P. Nassoy, and L. Cognet, “Self-interference 3d super-resolution microscopy for deep tissue investigations,” Nature Methods, vol. 15, pp. 449–454, apr 2018.

[45] P. Jouchet, C. Cabriel, N. Bourg, M. Bardou, C. Poüs, E. Fort, and S. Lévêque-Fort, “Nanometric axial localization of single fluorescent molecules with modulated excitation,” Nature Photonics, vol. 15, pp. 297–304, jan 2021.

[46] V. Levi, Q. Ruan, K. Kis-Petikova, and E. Gratton, “Scanning FCS, a novel method for three-dimensional particle tracking,” Biochemical Society Transactions, vol. 31, pp. 997–1000, oct 2003.

[47] K. C. Gwosch, J. K. Pape, F. Balzarotti, P. Hoess, J. Ellenberg, J. Ries, and S. W. Hell, “MIN-FLUX nanoscopy delivers 3d multicolor nanometer resolution in cells,” Nature Methods, vol. 17, pp. 217–224, jan 2020.

[48] N. Bourg, C. Mayet, G. Dupuis, T. Barroca, P. Bon, S. Lécart, E. Fort, and S. Lévêque-Fort, “Direct optical nanoscopy with axially localized detection,” Nature Photonics, no. August, 2015.

[49] J. Deschamps, M. Mund, and J. Ries, “3D superresolution microscopy by supercritical angle detection,” Optics Express, vol. 22, p. 29081, nov 2014.

[50] J. C. Thiele, M. Jungblut, D. A. Helmerich, R. Tsukanov, A. Chizhik, A. I. Chizhik, M. J. Schnermann, M. Sauer, O. Nevskyi, and J. Enderlein, “Isotropic three-dimensional dual-color super-resolution microscopy with metal-induced energy transfer,” Science Advances, vol. 8, jun 2022.

[51] C. Cabriel, N. Bourg, P. Jouchet, G. Dupuis, C. Leterrier, A. Baron, M.-A. Badet-Denisot, B. Vauzeilles, E. Fort, and S. Lévêque-Fort, “Combining 3D single molecule localization strategies for reproducible bioimaging,” Nature Communications, vol. 10, apr 2019.

[52] Y. Shechtman, L. E. Weiss, A. S. Backer, S. J. Sahl, and W. E. Moerner, “Precise three-dimensional scan-free multiple-particle tracking over large axial ranges with tetrapod point spread functions,” Nano Letters, vol. 15, pp. 4194–4199, jun 2015.

[53] A. Aristov, B. Lelandais, E. Rensen, and C. Zimmer, “ZOLA-3d allows flexible 3d localization microscopy over an adjustable axial range,” Nature Communications, vol. 9, jun 2018.

[54] B. P. Bratton and J. W. Shaevitz, “Simple experimental methods for determining the apparent focal shift in a microscope system,” PLOS ONE, vol. 10, p. e0134616, aug 2015.

[55] H. J. Anderson, J. K. Sahoo, R. V. Ulijn, and M. J. Dalby, “Mesenchymal stem cell fate: Applying biomaterials for control of stem cell behavior,” Frontiers in Bioengineering and Biotechnology, vol. 4, may 2016.

[56] M. Théry, A. Pépin, E. Dressaire, Y. Chen, and M. Bornens, “Cell distribution of stress fibres in response to the geometry of the adhesive environment,” Cell Motility and the Cytoskeleton, vol. 63, pp. 341–355, June 2006.

[57] C. S. Chen, J. L. Alonso, E. Ostuni, G. M. Whitesides, and D. E. Ingber, “Cell shape provides global control of focal adhesion assembly,” Biochemical and Biophysical Research Communications, vol. 307, pp. 355–361, July 2003.

[58] A. Barlian and K. Vanya, “Nanotopography in directing osteogenic differentiation of mesenchymal stem cells: potency and future perspective,” Future Science OA, vol. 8, jan 2022.

[59] Shuqin Cao and Quan Yuan, “An update of nanotopographical surfaces in modulating stem cell fate: a narrative review,” Biomaterials translational, vol. 3, no. 1, pp. 55–64, 2022.

[60] J. Park, S. Bauer, K. A. Schlegel, F. W. Neukam, K. von der Mark, and P. Schmuki, “Tio2 nanotube surfaces: 15 nm—an optimal length scale of surface topography for cell adhesion and differentiation,” Small, vol. 5, pp. 666–671, Mar. 2009.

[61] Y.-H. Kang, B. Choi, C. Ahn, S. Oh, M. S. Lee, and E.-J. Jin, “Titanium oxide nanotube surface topography and microrna-488 contribute to modulating osteogenesis,” Journal of Nanomaterials, vol. 2014, pp. 1–8, 2014.

[62] P. Kanchanawong, G. Shtengel, A. M. Pasapera, E. B. Ramko, W. Davidson, H. F. Hess, and C. M. Waterman, “Nanoscale architecture of integrin-based cell adhesions Pakorn,” Nature, vol. 468, no. 7323, pp. 580–584, 2011.

[63] T. Orré, A. Joly, Z. Karatas, B. Kastberger, C. Cabriel, R. T. Böttcher, S. Lévêque-Fort, J.-B. Sibarita, R. Fässler, B. Wehrle-Haller, O. Rossier, and G. Giannone, “Molecular motion and tridimensional nanoscale localization of kindlin control integrin activation in focal adhesions,” Nature Communications, vol. 12, may 2021.

[64] S. van de Linde, A. Löschberger, T. Klein, M. Heidbreder, S. Wolter, M. Heilemann, and M. Sauer, “Direct stochastic optical reconstruction microscopy with standard fluorescent probes,” Nature protocols, vol. 6, pp. 991–1009, jul 2011.

[65] G. T. Dempsey, J. C. Vaughan, K. H. Chen, M. Bates, and X. Zhuang, “Evaluation of fluorophores for optimal performance in localization-based super-resolution imaging,” Nature Methods, vol. 8, pp. 1027–1036, Nov. 2011.

[66] A. von Diezmann, M. Y. Lee, M. D. Lew, and W. E. Moerner, “Correcting field-dependent aberrations with nanoscale accuracy in three-dimensional single-molecule localization microscopy,” Optica, vol. 2, p. 985, nov 2015.

[67] A. Bouissou, A. Proag, N. Bourg, K. Pingris, C. Cabriel, S. Balor, T. Mangeat, C. Thibault, C. Vieu, G. Dupuis, E. Fort, S. Lévêque-Fort, I. Maridonneau-Parini, and R. Poincloux, “Podosome force generation machinery: A local balance between protrusion at the core and traction at the ring,” ACS Nano, vol. 11, pp. 4028–4040, apr 2017.

[68] I. Izeddin, J. Boulanger, V. Racine, C. G. Specht, a. Kechkar, D. Nair, a. Triller, D. Choquet, M. Dahan, and J. B. Sibarita, “Wavelet analysis for single molecule localization microscopy.,” Optics express, vol. 20, pp. 2081–95, jan 2012.

